# Haplotype-resolved reference genomes of the sea turtle clade unveil ultra-syntenic genomes with hotspots of divergence

**DOI:** 10.1101/2025.03.26.644878

**Authors:** Larissa S. Arantes, Tom Brown, Diego De Panis, Scott D. Whiting, Erin L. LaCasella, Gabriella A. Carvajal, Adam Kennedy, Deana Edmunds, Blair P. Bentley, Jennifer Balacco, Conor Whelan, Nivesh Jain, Tatiana Tilley, Brian O’Toole, Patrick Traore, Erich D. Jarvis, Oliver Berry, Peter H. Dutton, Lisa M. Komoroske, Camila J. Mazzoni

**Affiliations:** Department of Evolutionary Genetics, Leibniz Institute for Zoo- and Wildlife Research (IZW), Berlin, Germany; Berlin Center for Genomics in Biodiversity Research (BeGenDiv), Berlin, Germany; Marine Science Program, Department of Biodiversity, Conservation and Attractions, Kensington, WA 6151, Australia; Marine Mammal and Turtle Division, Southwest Fisheries Science Center, National Marine Fisheries Service, National Oceanic and Atmospheric Administration, La Jolla, CA, United States; Department of Biological Sciences, Florida Atlantic University, FL 33431, Florida, USA; New England Aquarium Rescue and Rehabilitation Department, Quincy, MA, USA; New England Aquarium Animal Health Department, Quincy, MA, USA; Department of Biological Sciences, Smith College, Northampton MA 01060 USA; Vertebrate Genome Laboratory, The Rockefeller University, NY, USA; CSIRO Environomics Future Science Platform, Indian Ocean Marine Research Centre, Crawley, Western Australia, 6009, Australia; University of Massachusetts Amherst, Department of Environmental Conservation, Amherst, MA, USA

**Keywords:** Cheloniidae, Reference Genomes, Genetic diversity, Demography, Synteny

## Abstract

**Background:** Reference genomes for the entire sea turtle clade have the potential to reveal the genetic basis of traits driving the ecological and phenotypic diversity in these ancient and iconic marine species. Furthermore, these genomic resources can support conservation efforts and deepen our understanding of their unique evolution.

**Results:** We present haplotype-resolved, chromosome-level reference genomes and high-quality gene annotations for five sea turtle species. This completes the catalog of reference genomes of the entire sea turtle clade when combined with our previously published reference genomes. Our analysis reveals remarkable genome synteny and collinearity across all species, despite the clade’s origin dating back more than 60 million years. Regions of high interspecific genetic distance and intraspecific genetic diversity are consistently clustered in genomic hotspots, which are enriched with genes coding for immune response proteins, olfactory receptors, zinc fingers, and G-protein-coupled receptors. These hotspot regions may offer insights into the genetic mechanisms driving phenotypic divergence among species, and represent areas of significant adaptive potential. Ancient demographic analysis revealed a synchronous population expansion among sea turtle species during the Pleistocene, with varying magnitudes of demographic change, likely shaped by their diverse ecological adaptations, and biogeographic contexts.

**Conclusions:** Our work provides genomic resources for exploring genetic diversity, evolutionary adaptations, and demographic histories of sea turtles. We outline genomic regions with increased diversity, linked to immune response, sensory evolution, and adaptation to varying environments that have historically been subject to strong diversifying selection, and likely will underpin sea turtle’s responses to future environmental change. These reference genomes can assist conservation by providing insights into the demographic and evolutionary processes that sustain and threaten these iconic species.

## 1. Background

The rapid loss of biodiversity, driven by erosion and destruction of habitats globally, underscores the urgent need to develop strategies to mitigate this crisis and safeguard the planet’s ecological balance, reversing declines in biodiversity. One of the fastest growing technologies for understanding biodiversity and supporting its management is genomics [1]. Recent advances in high-quality genomic resources have facilitated our abilities to explore the genetic underpinnings of Earth’s biodiversity, enabling a deeper understanding of the evolutionary and functional complexities of life. Initiatives such as the Earth Biogenome Project [2], European Reference Genome Atlas [3], Darwin Tree of Life [4] and Vertebrate Genomes Project [5] have driven standards and recommendations for the production of high-quality reference genomes for conservation of biodiversity. These initiatives have resulted in an ever growing database of high-quality reference genomes, which is expanding rapidly as technologies evolve.

This growth in genomic resources has allowed researchers to investigate the genetic bases of a number of features key to assisting in species management and conservation such as age and lifespan [6,7], sex [8], abundance [9] and community composition [10] among others. Focussing on the evolutionary adaptations of iconic or umbrella species within ecosystems gives us the opportunity to efficiently monitor biodiversity and assess the health of these ecosystems [11,12].

Sea turtles have existed since non-bird dinosaurs were roaming the Earth [13] and hold critical ecological roles in both oceanic and coastal environments, but are at threat globally due to anthropogenic activities such as direct harvest, fisheries bycatch, habitat loss and climate change, among other risks [14–16]. At present, three of the seven extant sea turtle species have been classified under IUCN criteria as ‘endangered’ (*Chelonia mydas* [17]) or ‘critically endangered’ (*Eretmochelys imbricata* and *Lepidochelys kempii* [18,19]) and a further three (*Lepidochelys olivacea*, *Caretta caretta* and *Dermochelys coriacea* [20–22]) have been classified as ‘vulnerable’. Finally, while listed as ‘data deficient’ under the IUCN Red List, *Natator depressus* [23] has been classified as ‘vulnerable’ by the Australian government [24]. Extensive conservation efforts have led to positive outcomes for many populations [25], however effort and success have not been universal, with some populations still in decline [26].

Sea turtle species exist around the globe, inhabiting a remarkable diversity of ecological niches [27], spanning from deep cold-water oceanic divers like *D. coriacea* to range-restricted endemic species, such as *N. depressus* and *L. kempii* [28]. For other species, their habitats span the tropics and sub-tropics (*L. olivacea)* and broader, temperate and tropical ranges, such as *C. mydas*, *E. imbricata*, and *C. caretta*. Some sea turtles demonstrate dietary specializations (*D. coriacea* and *E. imbricata*), while others (e.g. *C. mydas* and *C. caretta)* display generalist omnivorous feeding habits [29]. The genomic bases for these traits remain unclear, however having access to high-quality genomic resources would allow more fine-level investigation into genetic drivers behind the capabilities of sea turtles to live in varying habitats and adapt to changing conditions in the Anthropocene [30–32], such as identifying genes under selection, or areas of adaptive potential, made easier through the use of high-quality reference genomes for each species [33].

At present, genomes are available for five of the seven extant sea turtle species, namely from *C. mydas* and *D. coriacea* [34], *C. caretta* [35]*, E. imbricata* [36] and *L. olivacea* [37]. Previous analyses in particular of the genomes of *C. mydas* and *D. coriacea* that represent the two extant sea turtle families *(Dermochelyidae and Cheloniidae)* have revealed a high degree of synteny and collinearity within this ancient clade [34,38]. Alongside this high level of apparent genomic conservation, small highly divergent genomic regions have also been observed between these two species, in particular in areas containing multi-copy gene families such as the Major Histocompatibility Complex (MHC) and olfactory receptors [34] as well as some rearrangement of genes potentially involved in temperature-dependent sex determination [38]. While these are clearly important genomic regions for understanding sea turtle adaptation and evolution, it is not clear if the differences between *C. mydas* and *D. coriacea* are species specific, or how well they characterize comparative patterns within the entire sea turtle clade.

In this study, we add to our previous reference genomes for *C. mydas* and *D. coriacea* [34], by producing high-quality genomes for the remaining five extant sea turtle species. Our genomes are assembled using highly accurate PacBio HiFi (High-Fidelity) and Chromatin-Conformation-Capture (Hi-C) sequencing, producing genomes with chromosomes phased into both parental haplotypes. This first full catalogue of sea turtle genomes now provides a unique opportunity to understand and investigate the evolution of sea turtles and contextualise their evolution among other turtles and tortoises, spanning hundreds of millions of years of evolution. We uncover high levels of genome-wide synteny across all Testudine genomes, with a notable pattern of genetic diversity and divergence within the sea turtle clade, intricately clustered within specific regions of specific chromosomes. These regions are enriched in immune-related genes, suggesting a role in the adaptive capabilities of these species. Furthermore, we performed demographic, genetic diversity, and inbreeding analyses to offer further insights to inform and guide conservation efforts. Thus, we demonstrate the power of high-quality genomes to uncover complex patterns of genetic diversity and adaptation that are vital for understanding species evolution and informing conservation strategies.

## 2. Data Description

### 2.1. Sequencing

For the five turtle species (*C. caretta*, *E. imbricata*, *L. olivacea*, *L. kempii*, *N. depressus*), we sequenced PacBio HiFi reads ranging from 35x to 60x coverage for each genome (Fig S1) and Hi-C sequences ranging from 47x to 129x coverage. We sequenced optical maps with N50 values ranging from 222 to 266 kbp and total DNA yields from 103 to 498 Gbp for long-range molecules of higher quality for 3 of the five species (*L. olivacea*, *C. caretta* and *E. imbricata* Table S1). These datasets are available via the European Nucleotide Archive (ENA) and National Centre for Biotechnology Information (see Data Availability).

### 2.2. Genome Assembly

Our haplotype-separated chromosome-scale assemblies are highly contiguous, and in particular, are significantly more contiguous than the previously published *D. coriacea* assembly based on PacBio CLR data and *C. caretta* assembly based on Oxford Nanopore Technology (ONT) reads (Figs 1a, S1 & S2, Table S2). Moreover, the new haplotype assemblies show exceptional base accuracy, with a Quality Value (QV) ranging from 65.2 to 70.4 (Table S2). For comparison, the older CLR-based assemblies display QVs in the range of 38.7 to 47.6, while the ONT-based is much lower (in part due to a different sample used for analysis, Table S2). All genomes were scaffolded into complete chromosome molecules, with between 99.1% and 99.9% of the assembled sequences assigned to the 28 chromosomes (Figs 1b, S3 & S4, Table S2). The assembled genomes also show high gene completeness, with between 98.5% and 99.5% single-copy orthologs from the Sauropsida lineage identified by BUSCO (Figs 1c & S5, Table S2).

**Figure 1.**
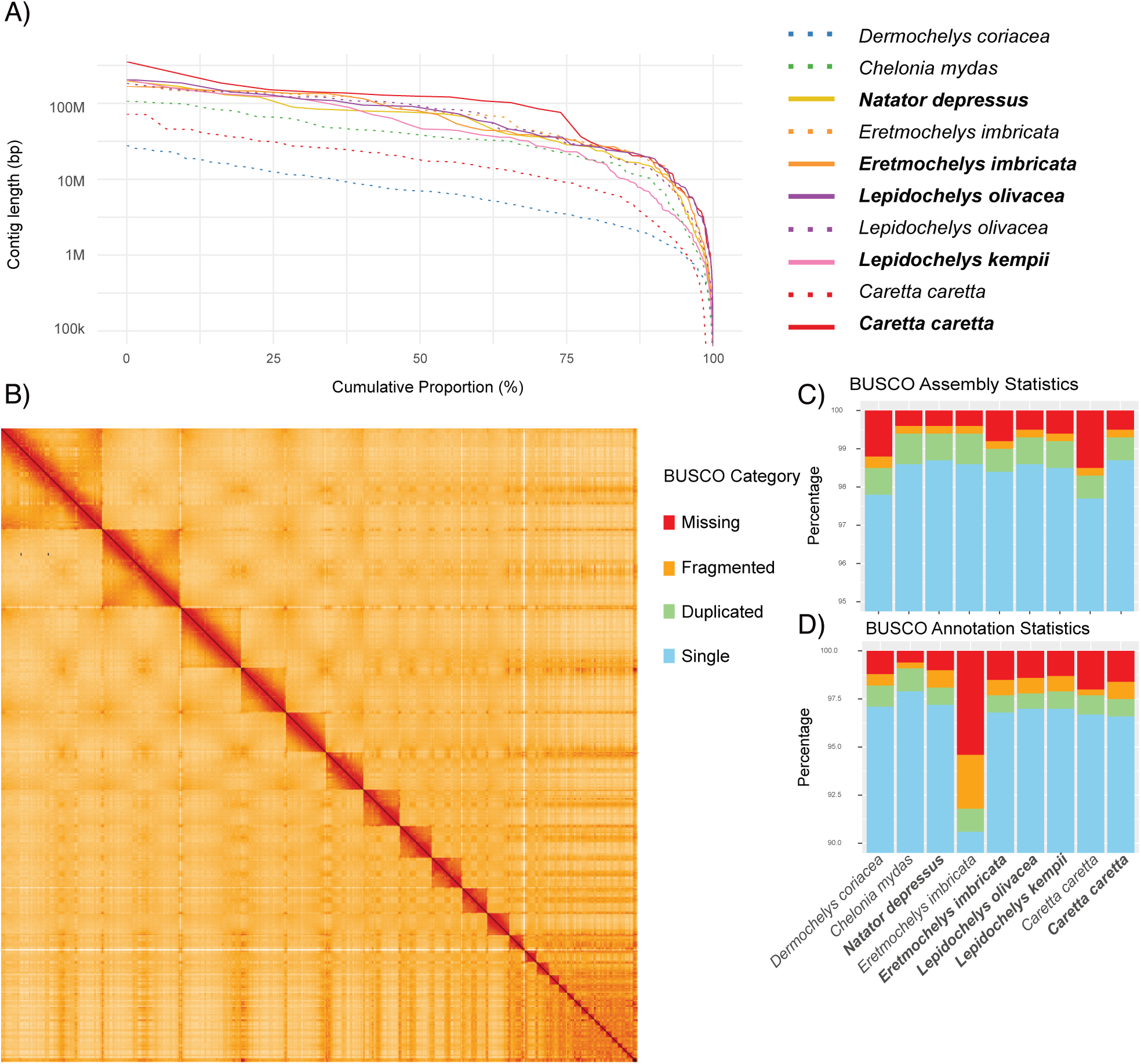
A) Lengths of the assembled contigs for each species sorted by length (y-axis) and scaled to total length of each genome (x-axis). Genomes from this study are shown as complete lines with names in bold, previously published assemblies are shown as dashed lines. B) 3-dimensional conformational arrangement of one assembled *Caretta caretta* haplotype genome assembly as evaluated by Hi-C. The x- and y-axes show the coordinates of the respective genome and each detected contact in the genome is coloured with increasing intensity from white to red. The red diagonal shows the self-interactions of each position with itself and close vicinity, squares show the high self-interaction of chromosomes. C) Percentage detected single copy orthologs identified in the genome assemblies calculated via BUSCO using the Sauropsida database. Scores are calculated based on detected mappings of ortholog sequences using the Miniprot mapper. D) Gene completeness of sea-turtle protein-coding annotations based on BUSCO genes identified in each annotated protein set.

### 2.3. Genome Annotation

Using a combination of approaches based on transcriptomic data, protein sequences from *C. mydas* and *D. coriacea*, liftover annotations from *C. mydas* and *Malaclemys terrapin pileata,* as well as *de-novo* gene predictions, we created a set of protein-coding gene predictions for each of our assembled genomes (Table S3). The annotations themselves are highly complete when evaluated based on single-copy orthologs from Sauropsida via BUSCO (Fig 1d) and hierarchical orthology groups from Archelosauria via OMArk (Fig S6), reaching comparable completeness to previous annotations generated by RefSeq, with BUSCO scores between 97.2% and 98.1% and OMArk completeness scores between 97.46% and 98.47%, furthermore capturing many BUSCO genes missing in the annotation provided for the existing *E. imbricata* reference genome [36].

## 3 Analyses

### 3.1 Genome Synteny

Based on identification of orthologous genes and their locations in Testudine genomes, we uncovered remarkably high synteny across the clade, encompassing over 100 million years of evolution with only a small number of hotspots of variation identified. Particularly among the sea turtles, all 28 chromosomes were highly collinear and syntenic (Figs 2a & S7) with complete one-to-one synteny demonstrated at the chromosome level, with the exception of one region at the end of chromosome 14 in *D. coriacea*, found in chromosome 11 in the six *Cheloniidae* turtle species (Fig S7).

**Figure 2.**
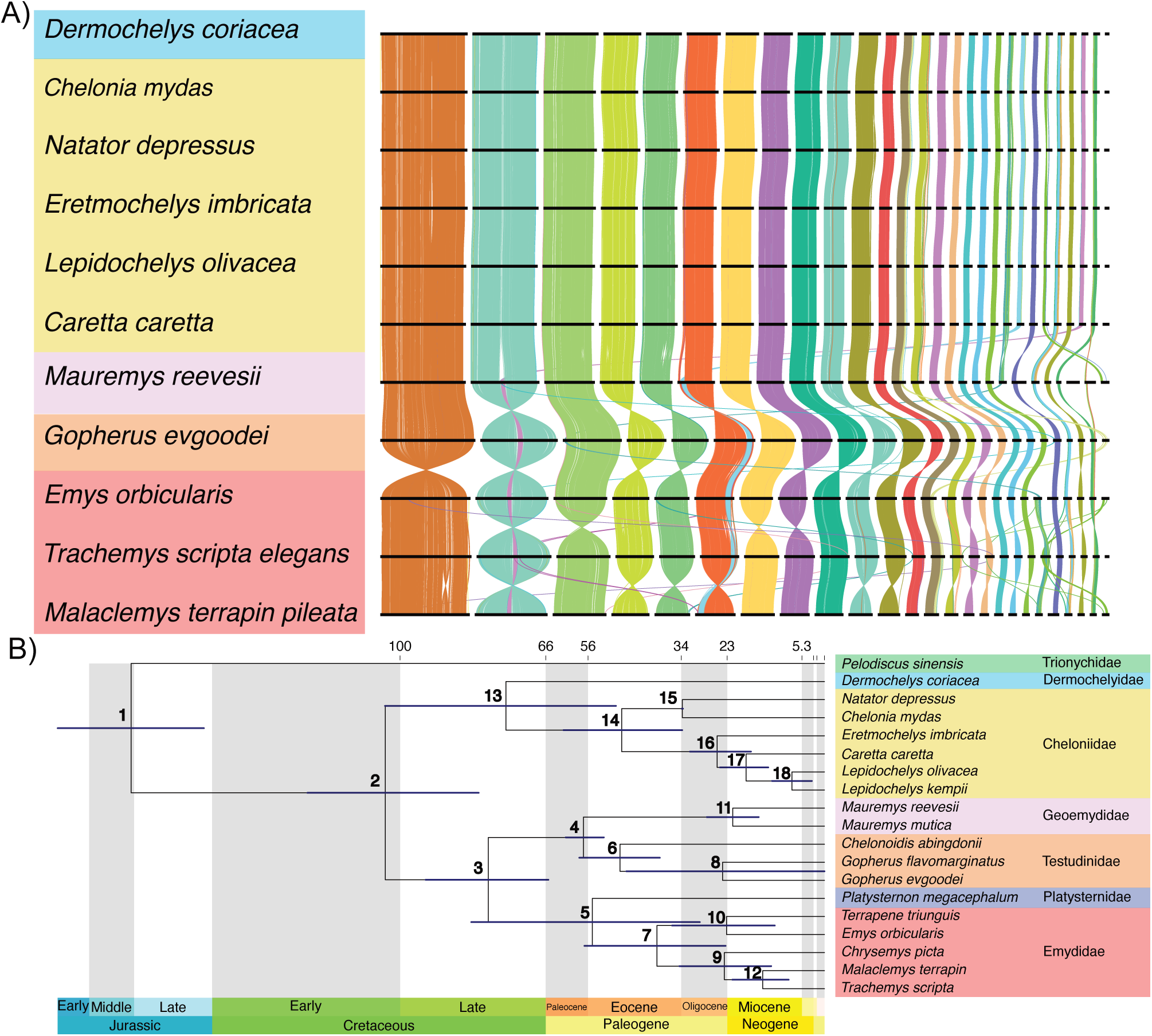
A) Genome-wide gene-synteny plots across Testudines. Each line represents a best-reciprocal-hit protein match between annotated genes in each consecutive genome. Lines are coloured based on co-localisation across all 11 genomes determined by Fisher’s Exact Test. Chromosomes are ordered based on synteny to Dermochelys coriacea. B) Divergence times of species within the suborder Cryptodira based on protein-coding genome annotations. The bars on each node represent the 95% highest posterior density (HPD) intervals for node age estimates. Divergence times and confidence intervals for each numbered node are detailed in Table S4.

Across Testudine genomes (i.e. including terrestrial and freshwater turtle and tortoise families; Fig 2), we found that the macrochromosomes (>50Mb in length) exhibited high synteny across all turtle genomes and among the microchromosomes (<50Mb in length), with only chromosomes 21 and 26 from sea turtles rearranged in other turtle genomes. In these instances, chromosomes 21 and 26 from the sea turtle genomes were found in the arm of chromosome 4 (which is syntenic to chromosome 6 in the sea turtle genomes), and the central region of chromosome 2, respectively, in the genome of the Chinese pond turtle (*Mauremys reevesii)* with this positioning conserved across all other Testudine genomes (Figs 2a & S8).

### 3.2 Phylogenomic analysis

Phylogenetic analysis using coding-protein sequences for all turtle species with annotation available provided insights into evolutionary relationships and speciation events within suborder Cryptodira, which includes most living turtles and tortoises (Fig 2b). The topology and divergence time support the findings of previous studies based on a few nuclear markers or mitochondrial DNA [39–41] (Table S4). Our genome-wide analysis indicates that the sea turtle clade diverged from other Durocryptodira species 104 million years ago (mya) [95% highest posterior density (HPD) = 81.9 to 122 mya]. Dermochelyidae (including *D. coriacea*) separated from the Cheloniidae family approximately 75.4 mya (95% HPD = 49.4, 104). Within the Cheloniidae family, the divergence of *C. mydas* and *N. depressus* occurred approximately 33.6 mya (95% HPD = 33.5, 33.8), while the other species diverged around 25.4 mya (95% HPD = 17.4, 31.9). *Lepidochelys kempii* and *L. olivacea* were the most recently diverged lineages, having split around 7.72 mya (95% HPD = 2.99, 12.4), a time period associated with significant environmental changes such as the closure of the Tethys Sea and cooling of the southern oceans, which likely disrupted gene flow and contributed to the speciation of these two *Lepidochelys* species [40,42].

### 3.3 Genome-wide patterns of diversity and divergence

Comparisons of within-individual genetic diversity, measured by average heterozygosity per chromosome, revealed consistent variation across chromosomes in the individuals sequenced, with each species showing distinct magnitude of variation (Figs 3a & S9). Notably, chromosomes with high intraspecific diversity also exhibited higher gene density (Fig 3b) and increased interspecific genetic distance (Fig 3c). Consistent with findings in sea turtles and other species with microchromosomes [43], we found that the average heterozygosity, as well as the gene density and interspecific genetic distance, were higher for microchromosomes (12-28) than macrochromosomes (1-11) (p<0.05) (Figs 3a-c). In particular, chromosomes 13, 14, 20, 23, 24, and 28 exhibited heightened genetic diversity, interspecific divergence, and gene density (Figs 3a, 3c, S9 & S10, Table S5).

**Figure 3.**
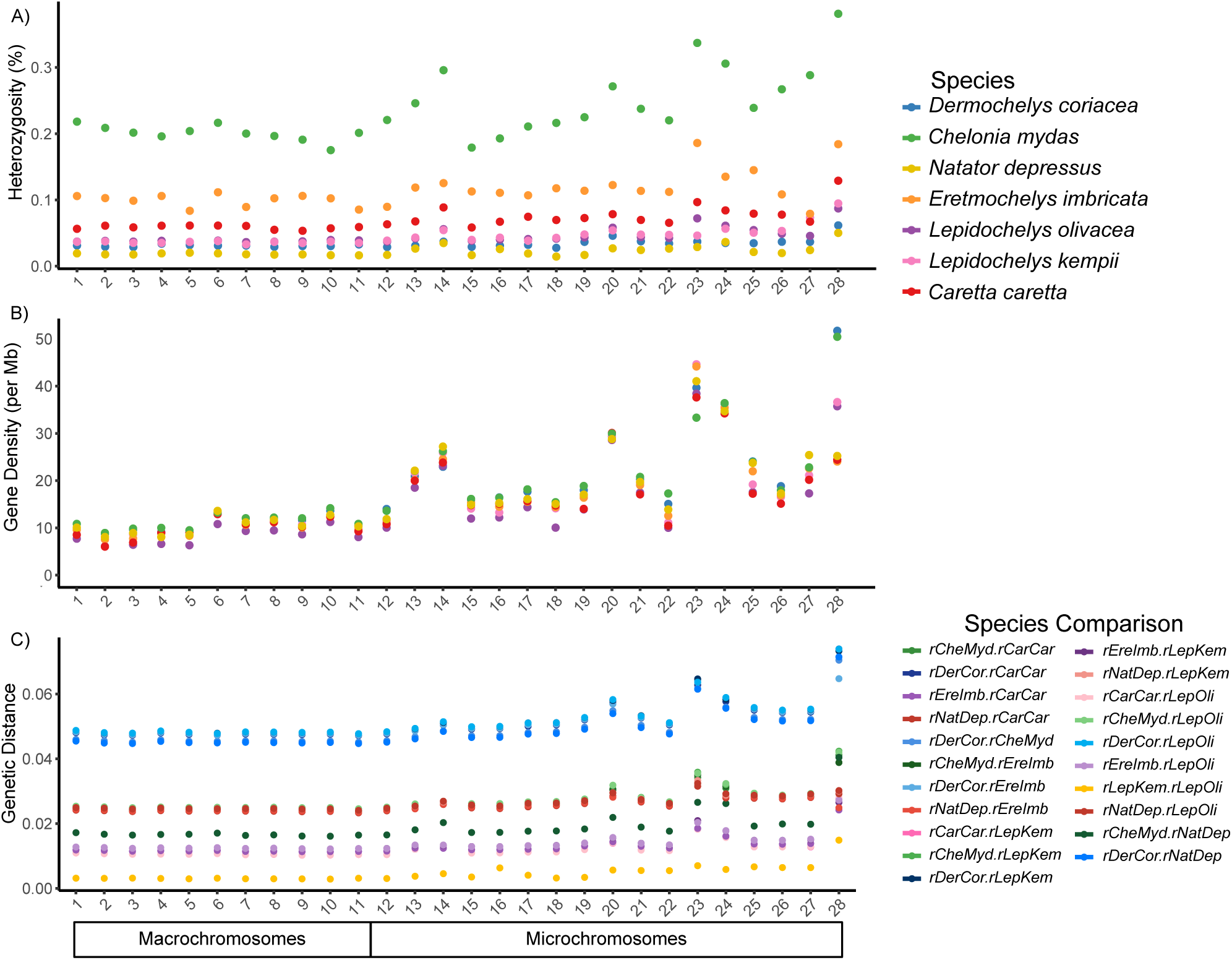
A) Heterozygosity, B) gene density (per Mb) and C) pairwise genetic distance per chromosome for the seven sea turtle reference genomes. Chromosomes longer or shorter than 50 Mb are highlighted as macrochromosomes or microchromosomes, respectively.

Furthermore, increased levels of heterozygosity and interspecific genetic distance were concentrated at particular hotspot regions, defined as regions with heterozygosity exceeding four times the chromosomal mean and genetic distance double that of the chromosomal mean, rather than being uniformly distributed across an entire chromosome (Fig S9). Thus, we identified the three regions located in chromosomes 13, 14 and 24 exhibiting colocalised elevations in heterozygosity and genetic distance across sea turtles (Fig 4). While we identified these hotspot regions by calculating genetic distances from all species in relation to *D. coriacea* (Fig 4b), this pattern is consistent across pairwise comparisons between all species (Figs S10 & S11).

**Figure 4.**
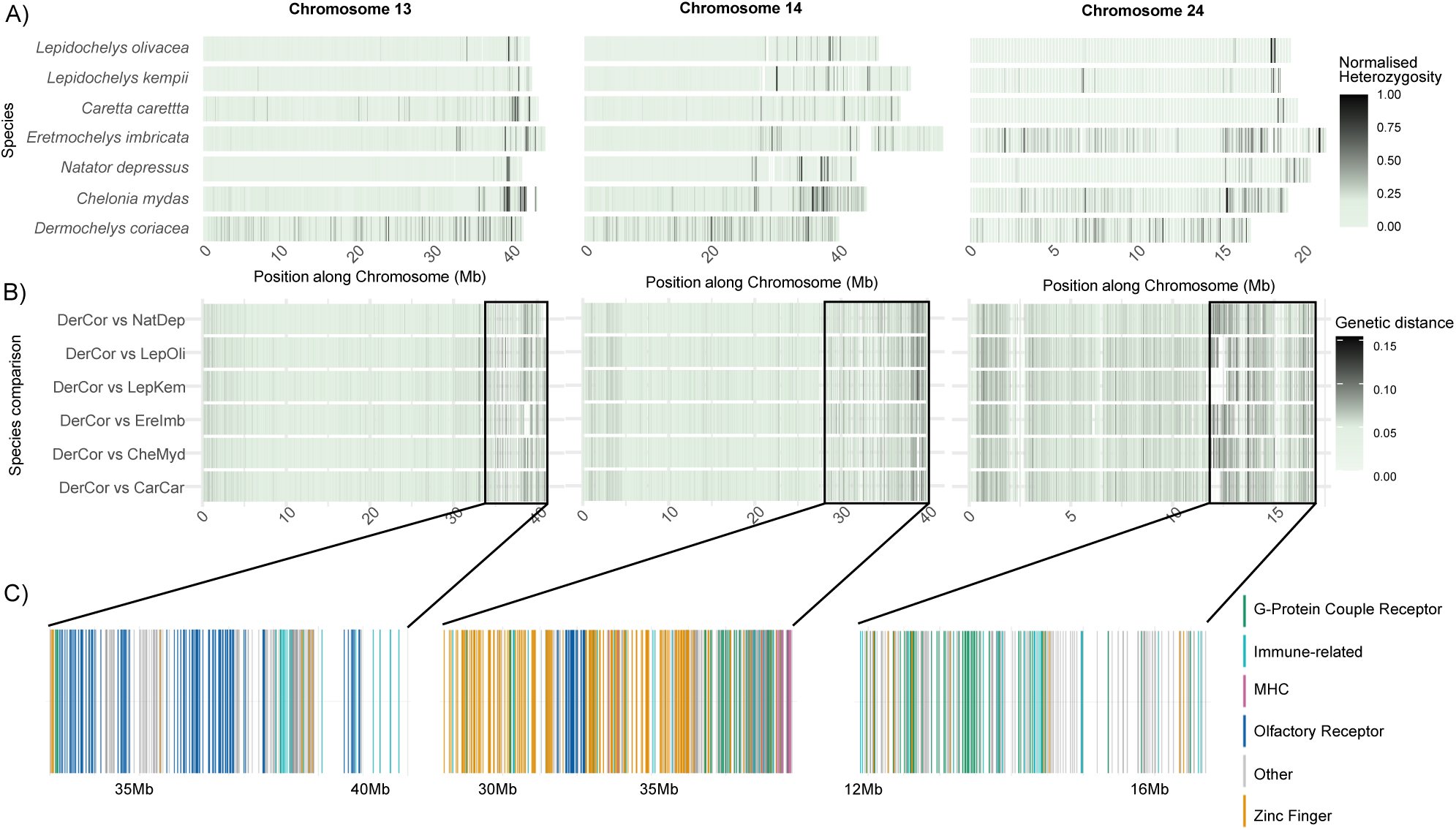
Genetic diversity and divergence hotspots contain genes associated with immune response, olfactory receptors, zinc fingers, and G-protein-coupled receptors. A) The heatmap illustrates normalised heterozygosity (He) across chromosomes 13, 14, and 24 for seven sea turtle species, displaying He values in non-overlapping 50 kb windows. The normalisation highlights chromosomal hotspots rather than overall diversity. B) Pairwise genetic distances between the six sea turtle species and D. coriacea are shown along the same three chromosomes. Genetic distance was calculated as the ratio of interspecific single variants per 10 kb. Black boxes highlight the chromosome areas of increased genetic distance among sea turtle genomes. C) Multi-copy gene families located in the highlighted regions are displayed and colour-coded by their annotation. Chromosome coordinates are shown by their position in the genome of *D. coriacea*.

Following functional annotation of the genes found in these hotspots, we found enrichment for multi-copy gene families coding for proteins with functions in immune response, olfactory receptors (ORs), zinc fingers, and G-protein-coupled receptors (GPCRs_ (Fig 4c, Tables S6 & S7). This included enrichment of immunology-related genes, GPCRs, ORs, and Zinc-finger genes in chromosome 13 (adjusted p < 10^-42^, 10^-47^, 10^-79^, 0.01, respectively), MHC genes, Immunology-related genes, GPCRs, ORs, and Zinc-finger genes in chromosome 14 (adjusted p < 10^-24^, 10^-6^, 10^-2^, 10^-10^, 10^-52,^ respectively) and Immunology-related genes and GPCRs in chromosome 24 (adjusted p < 10^-3^ and 10^-3^, respectively). A particular concentration of olfactory receptors - known for their role in odor perception and detection of chemical cues, was identified in the hotspot region of chromosome 13 (Fig 4) and Major histocompatibility complex (MHC) genes were concentrated within the identified hotspot on chromosome 14.

### 3.4 Inbreeding and Demographic histories

To investigate the levels of historical inbreeding within each species, we measured the proportion of the genome composed of runs of homozygosity (FRoH). The *N. depressus* individual sequenced in this study had the highest level of inbreeding (FRoH = 0.227), while *L. olivacea* showed the lowest (FRoH = 0.0149) (Fig 5a). The reference *C. mydas* individual had the third-highest inbreeding level (Fig S9). This is consistent with the observations made by Bentley et al [34], with this individual originating from a small breeding population in the Mediterranean sea with maternal and paternal lineages likely sharing a common ancestor in the relatively recent past.

**Figure 5.**
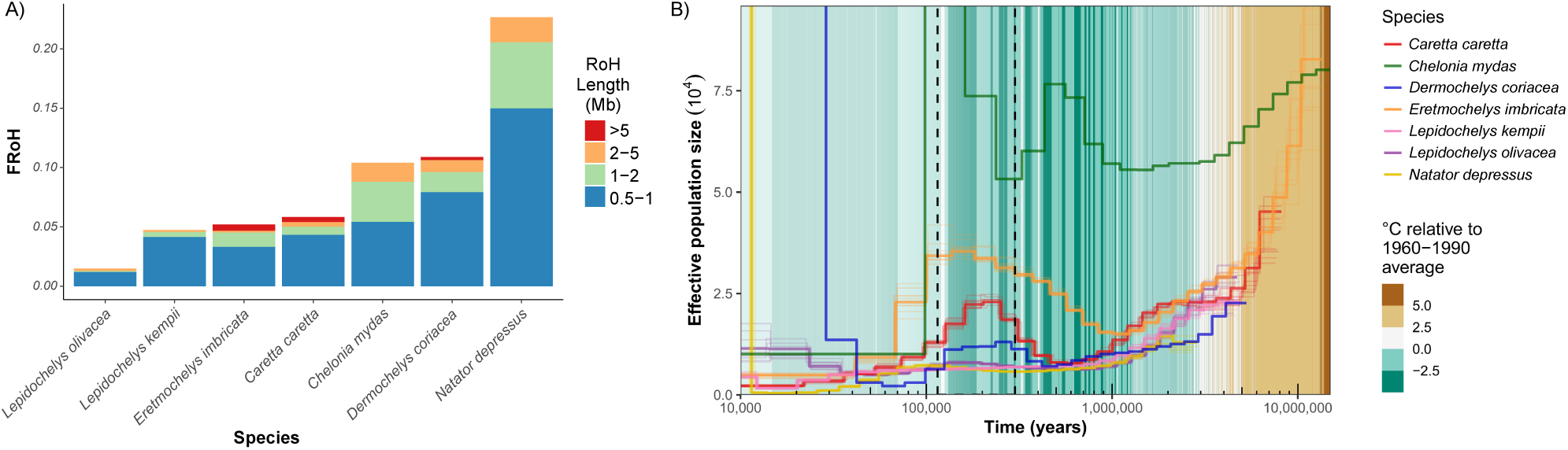
A) Inbreeding levels for the seven sea turtle individuals, measured as the proportion of the genome in runs of homozygosity (FRoH). FRoH are categorised by length (in Mb), where longer runs indicate more recent events associated with a shared common ancestor of the individual’s maternal and paternal lineages, while shorter runs suggest older inbreeding events. B) Ancient demographic history for the seven sea turtle species reconstructed with Pairwise Sequential Markovian Coalescent (PSMC) plot. Dashed lines indicate the Last Interglacial (Eemian Period, 130,000 to 115,000 years ago). Bootstrap replicates (10 for each lineage) are plotted in lighter lines. Inferred fluctuations in effective population size (Ne) were rescaled assuming 30-year generation time and 1.2 × 10−8 per generation mutation rate.

Using Pairwise Sequentially Markovian Coalescent (PSMC) models [44], we reconstructed the demographic histories of the seven extant sea turtle species, revealing consistent patterns of population declines beginning approximately 1–9 mya, likely driven by cooler sea temperatures (Fig 5b). During the Pleistocene, between 400 kya and 1 mya, all species, except *L. kempii*, experienced synchronous population expansions, with the growth particularly pronounced in *C. mydas*, *C. caretta* and *E. imbricata*, while *L. kempii* maintained a relatively stable population size. The population peak occurred during the Last Interglacial in the Eemian period (115-130 kya). This period of growth was followed by a second population decline across all species, starting roughly 100 kya until recently around 50 kya. The three species with relatively stable historical population sizes - *N. depressus*, *L. kempii*, and *L. olivacea* - differ significantly in their levels of genetic diversity. *N. depressus* exhibits the lowest heterozygosity, while the two *Lepidochelys* species display relatively high heterozygosity.

## 4. Discussion

Our chromosome-scale, annotated genomes across the sea turtle clade revealed remarkable genetic synteny across this slowly-evolving group of animals, while also revealing hotspot regions of the genome consistently undergoing accelerated evolution and divergence that likely play important roles in the morphological and ecological diversity exhibited among these species. These regions contained genes important for immune responses, the ability to sense and respond to the environment and regulate gene expression under fluctuating environmental conditions [45]. This builds on previous results comparing genomes of *C. mydas* and *D. coriacea* [34], demonstrating that sea turtle genomes have remained highly syntenic since their split from freshwater turtles and tortoises around 100 million years ago.

The availability of high-quality reference genomes for all sea turtles opens new avenues to explore fundamental questions about their adaptation, immunity, and sensory evolution. Sea turtles exhibit remarkable adaptations to marine environments, including extreme migratory behaviors [46], saltwater tolerance [47], natal homing [48,49], and temperature-dependent sex determination [50], yet the genetic basis of these traits remains poorly understood. Our results highlight microchromosomes and specific regions of reduced relative synteny in macrochromosomes as key loci enriched in gene density and genetic variation across the sea turtle clade. This pattern is also observed in birds and other reptiles, with the high GC content and high recombination rate of the microchromosomes potentially playing a significant role in promoting diversification [43].

The highlighted hotspots of evolutionary diversification harbor multicopy gene families, such as olfactory receptors involved in detecting odorants and adapting to the chemical complexity of habitats [51,52], as well as MHC genes, central to the immune response to diseases [53,54]. These multicopy gene families found within divergent hotspots may represent adaptation mechanisms that maintain flexibility in response to dynamic or disruptive selective pressures, potentially aiding immune variability, environmental sensing, and essential survival responses across the diverse habitats these turtles inhabit. Sea turtles are known to inhabit a vast proportion of the globe’s seas, found in both deep and shallow waters [47,48], migrating long distances across highly variable temperatures, currents and salinity [37], as well as possessing an immune system highly influenced by these changing environments [49,50]. As enhanced MHC variation is associated with lower disease susceptibility [55], the MHC gene copy numbers and heterozygosity in sea turtles have been previously proposed to vary among species based on their habitats, with those exposed to higher pathogen loads and diversity in neritic environments exhibiting greater MHC gene copy numbers than species inhabiting pelagic habitats, an area which would require manual validation in future studies [34]. Thus, these genome hotspots of increased diversity and divergence may hold the key to understanding chemosensory evolution, disease resistance, and phenotypic diversity in sea turtles. Further exploration of these regions could shed light on adaptive forces that have influenced the evolutionary trajectory of sea turtle species.

From a conservation perspective, genomic resources offer powerful tools to help understand sea turtle population viability and resilience to anthropogenic threats. Genomic diversity, inbreeding levels, effective population sizes, and demographic histories are critical metrics for assessing extinction risk and adaptive potential [56]. Our results indicate a long-term trend of lower population sizes and genetic diversity for *N. depressus*, similar to the demographic trajectory observed for *D. coriacea* [34], rather than a sharp loss due to recent declines, highlighting the need to distinguish historical demographic patterns from contemporary inbreeding. While reduced diversity may have been sustainable in the past, potentially leading to some degree of purging of deleterious alleles, it could still limit adaptive capacity in the face of rapid environmental change [57]. In contrast, *Lepidochelys* species exhibit comparatively high genetic diversity despite their historically small population sizes, with *L. kempii* retaining higher genetic variation even with its restricted distribution in the Gulf of Mexico (Table S5). This contrasts with the other range-restricted sea turtle species, *N. depressus*, suggesting that endemism alone does not consistently predict genetic diversity in sea turtles. We acknowledge that our results are based on a single individual, and the individual’s origin should be considered, as previous studies have highlighted different demographic histories and genetic diversity between ocean basins [58]. However we have previously shown demographic histories of *C. mydas* and *D. coriacea* to be consistent even from individuals in different populations [34]. In the case of range-restricted species such as *N. depressus* and *L. kempii*, we anticipate that the demographic histories likely reflect range-wide patterns.

The demographic histories of sea turtle species reveal broadly similar trajectories of expanding and contracting effective population size over the past ten million years, though with the magnitude of N_e_ varying between the species. These unique patterns likely reflect the intersections of species-specific life histories and changing environments such as fluctuations in ocean temperature, sea level, and connectivity [59,60]. Species inhabiting shallow coastal habitats, such as *N. depressus*, were likely particularly affected by the dynamic coastal environment [61]. Specifically, the low and stable population size of *N. depressus* may reflect its restricted neritic distribution and tendency to disperse over smaller distances compared to other sea turtle species, potentially limiting foraging opportunities, preventing the species from achieving the global distribution exhibited at some other turtles [62]. On the other hand, the historically low population size of *D. coriacea* may be attributed to its specialized open-ocean cold-water lifestyle and high trophic position, primarily consuming gelatinous zooplankton, along with behavioral constraints tied to its large size and the challenges of terrestrial nesting [27]. The early pleistocene glaciation appears to have impacted dermochelyids more severely than the chelonids, resulting in the extinction of all but one of the dermochelyid species [64], which subsequently entered the Pleistocene expansion as a severely bottlenecked remnant population [63]. Conversely, *E. imbricata*, *C. caretta*, and *C. mydas*, fared better during the glacial contraction [64] and experienced more pronounced population expansions during the Pleistocene. The ability of *E. imbricata* to exploit diverse habitats and food sources, with a diet centered on coral reef organisms, likely favored its population expansion by reducing interspecies competition [65]. Similarly, *C. caretta* and *C. mydas* may have benefited from their broad dietary flexibility and ability to thrive in diverse temperate and tropical environments [27].

We observed a strong synchrony in population expansions, with population peaks during the Eemian period (130,000-115,000 years ago), although the magnitude of demographic changes varied among lineages. Reid et al. (2019) [58] also reported a synchronized demographic response after the Last Glacial Maximum across most sea turtle lineages. These population expansions most likely helped maintain genetic diversity in these species. Expanding these comparisons to include individuals from additional populations might further corroborate the links between demographic history and ecological factors such as habitat specificity, feeding habits, thermal preference, developmental and adult foraging stages (oceanic vs. neritic), and environmental conditions. This will be particularly valuable for estimating recent changes in population size, which rely on population-level genomic resources [66], and for understanding their connection to human-mediated environmental disturbances.

## 5. Potential Implications

High-quality reference genomes are important building blocks for creating genomic toolkits for species conservation and management. One exciting consequence of discovering the levels of genome-wide synteny exhibited between sea turtles is that genetic markers identified for determining features such as sex and adaptive traits in one species may also be directly applicable to other species without the need for new rounds of research and development. Having complete, annotated, chromosome-level genomes for all sea turtles means that such markers or genetic regions can be quickly verified between the species and turned into practical conservation toolkits. While they may not be required for individual studies with a scope of a single or few populations, they are critical for anchoring markers and comparing across studies and species. For example, the advancement from mitochondrial to whole-genome markers help alleviate conflicting signals that can arise from nuclear integrations of mitochondrial sequences (NuMTs), recently misinterpreted as evidence of a new species of *D. coriacea* [67,68], giving better resolution to future genomic studies with potential conservation implications. In this case the identified NuMT contained a portion of the mtDNA Control Region, commonly used for population structure analysis in all the sea turtle species, however we did not find such NuMTs in the other genomes reported here.

We believe these reference genomes will also be valuable for measuring and predicting the impact of climate change on sea turtles. For ectotherms like reptiles, climate impacts may be particularly pronounced due to their sensitivity to thermal fluctuations [69]. For sea turtles, these effects have potential to be even greater due to their temperature-dependent sex determination, where changes to nest temperature can disrupt sex ratios and reproductive success [50,70]. As these effects may be best evidenced via the epigenome, having access to complete, annotated reference genomes increases the predictive power of markers based on measuring levels of DNA or chromatin modifications.

Our findings highlight how different sea turtle species have responded to ancient climate changes, reflecting a range of adaptive strategies and unique biogeographic scenarios. Understanding how species have historically responded to changes in climate offers insights into their potential reactions to current and future anthropogenic disturbances, helping to inform conservation strategies and predict the long-term impacts of climate shifts on sea turtle populations.

## 6. Methods

### 6.1. Sampling

Whole blood samples were collected from juvenile *C. caretta, L. kempii, L. olivacea* and *E. imbricata,* and immediately flash-frozen at -80°C. A blood sample from a female *N. depressus* was stored in ice for 24 hours before being frozen at -80°C. Additionally, organ tissue samples were collected opportunistically from *L. kempii* (brain, kidney, and ovary) and *C. caretta* (thymus, ovary, brain, liver, heart, spleen, testes, kidney, and lung) and flash frozen at -80°C for long and short read transcriptomic sequencing for genome annotation. We shipped the samples on dry ice or in liquid nitrogen dry shipper, ensuring that they remained consistently frozen throughout transit.

### 6.2. Sample Processing and Sequencing

We extracted and purified DNA using a Bionano SP DNA kit (PN 80042) for *C. caretta*, *E. imbricata*, and *L. kempii*. We used a MagAttract HMW DNA Kit (Qiagen 67563) for *N. depressus* and *L. kempii*. We measured DNA quantity using triplicate measures and Qubit 3 fluorometer (Invitrogen Qubit dsDNA Broad Range Assay cat no. Q32850) and measured DNA size with an Agilent Femto Pulse. We fragmented the DNA to 15 – 20 kb length prior to library preparation using a Megaruptor 3 (Diagenode, Denville, NJ, USA) and standard hydropores (Cat. No. E07010003).

We prepared the PacBio HiFi libraries using a SMRTbell prep kit 3.0 (Pacific Biosciences PN 102-182-700) and PacBio barcoded primers. We size-selected the libraries to remove DNA under 10kb using a Pippin HT instrument (Sage Science, Beverly, MA, USA). We then quantified the size-selected HiFi libraries with a Qubit 3 Fluorometer (Qubit dsDNA HS Assay Kit), and assessed the average size with an Agilent Femto Pulse.

For *C. caretta*, *E. imbricata*, and *L. kempii* we sequenced HiFi libraries with a PacBio Sequel IIe instrument on 8M SMRT cells (101-389-001) using Binding kit 3.2 (102-333-300) and Sequel II sequencing kit 2.0 (101-820-200), and 40-hour movie time with 2-hour pre-extension. For *N. depressus* and *L. olivacea* we sequenced HiFi libraries with a PacBio Revio instrument using a Revio polymerase kit (102-817-600), Revio sequencing plate (102-587-400), and 24-hour movie with 1.6-hour pre-extension.

For *C. caretta*, *E. imbricata*, *L. kempii* and *L. olivacea*, we prepared Omni-C libraries using the Dovetail Omni-C Kit (Dovetail Genomics, CA) according to the manufacturer’s protocol. We then sequenced the Omni-C libraries with the Illumina NovaSeq 6000 platform with 2×150 bp read length. For *N. depressus* we prepared the Hi-C library using the Arima-HiC 2.0 kit (Arima Genomics, Carlsbad, CA, USA) following the manufacturer’s protocol. We then sequenced the Hi-C libraries with the Illumina NovaSeq 6000 platform with 2×150 bp read length.

For Bionano optical mapping, we labelled 750 ng DNA using direct labeling enzyme (DLE1) and the Bionano Prep Direct Label and Stain (DLS) protocol (document number 30206) and then imaged the DNA on the Bionano Saphyr instrument.

To prepare RNA for sequencing, we extracted and purified total RNA using a QIAGEN RNAeasy kit (cat. 74104). We determined the RNA quantity using a Qubit 3 fluorometer (Invitrogen Qubit RNA High Sensitivity (HS) Kit (cat. no. Q32852)) and measured the RNA integrity (RIN) score using an Agilent Fragment Analyzer. We prepared the RNA-Seq libraries using the Illumina Stranded mRNA Prep kit and sequenced the libraries with the Illumina NovaSeq 6000 platform with 2×100bp read length. We generated Iso-Seq cDNA libraries using the NEBNext Single Cell/Low Input cDNA Synthesis & Amplification Module in combination with PacBio’s SMRTbell Prep Kit 3.0. We then sequenced the Iso-Seq libraries on a PacBio Sequel IIe machine using a Sequel II 8M SMRTcell.

### 6.3. Genome Assembly

We performed the assemblies of each genome following the best-practices established by the Vertebrate Genomes Project [5,71]. In particular, we trimmed the raw sequencing reads for adapters using cutadapt v4.9 to remove any remaining PacBio adapter sequences from the PacBio HiFi reads and Illumina adapters from the Hi-C reads. We assembled initial contig sets for each species using hifiasm [72], v0.19.4-9, l2-l3, Hi-C phasing mode, using both PacBio HiFi and Illumina Hi-C reads as input to generate two haplotype-phased sets of contigs. We then removed retained haplotigs from each assembly with purge-dups [73] v1.2.6, -e. To scaffold the assembled contigs into chromosomes, we used the hybrid-scaffold tool from the Bionano Solve suite (v3.7.0, VGP mode) to scaffold with optical maps and then mapped the Hi-C reads to the set of initial scaffolds using bwa-mem [74] v2.2.1, -5SP -T0 and scaffolded into pseudo-chromosomal units using yahs [75] v1.2a.1. Finally, we performed rounds of manual curation following the Sanger rapid-curation pipeline [76], joining any missed-scaffolds and removing any false joins in the assembly. We screened for any retained adapter or vector sequences using ncbi’s FCS-adapter and for foreign contaminant sequences using ncbi’s FCS-GX [77] v0.5.4.

### 6.4. Genome Annotation

To generate a set of protein-coding annotations for each genome, we incorporated evidences from transcript data, protein sequences, *ab-initio* machine-learning approaches and homology to genomes of related species. To create *ab-initio* predictions, we ran Helixer [78] vv0.3.3_cuda_11.8.0 using argument *–lineage vertebrate*. To generate protein-based gene model predictions, we mapped protein sequences from existing *C. mydas* and *D. coriacea* assemblies (GCF_015237465.2 and GCF_009764565.3, respectively) using miniprot [79] v0.13-r248. To create transcript-based gene model predictions, we mapped paired-end RNA-seq data to each genome using hisat2 [80] v2.2.1 using argument *–dta* and filtered the alignments using samtools [81] v1.19.2 with argument *-F 3840*. We then generated a *de-novo* transcript assembly using stringtie [82] v2.2.1 and predicted coding sequences using TransDecoder [83] v5.7.1 and included only those gene models with a TransDecoder score greater than 20. Similarly, we mapped PacBio Iso-seq data to the genome using minimap2 [84] v2.28-r1209 with argument *-x splice:hq* and filtered the alignments using samtools with argument *-F 3840* and built gene models using stringtie with argument *-L* and predicted CDS using TransDecoder as above. To generate homology-based predictions, we created lastz-alignment chains from *C. mydas* and *Malaclemys terrapin pileata* genomes (GCF_015237465.2 and GCF_027887155.1) using the *make_lastz_chains* (v2.0.8) tools from TOGA [85] and we generated the set of homology gene predictions using TOGA (v1.1.6).

To generate a set of best gene models, we used EvidenceModeler [86] v2.1.0 to combine all of the above evidences using the weights defined in Table S8.

To create functional annotations, we mapped the amino acid sequences from each gene model against the swissprot database [87], release 2023_03 using the diamond [88] v2.1.8 blastp search and we identified Pfam, PROSITE and SUPERFAMILY homology using Interproscan [89,90] v5.59-91.0. Finally, we filtered gene models which had no identified swissprot or Pfam homology and were over 50% masked, or missing start and/or stop codons.

### 6.5. Genome synteny

To determine the number and sizes of syntenic regions within turtle and tortoise genomes, we made use of the annotated protein sequences to find orthologous genes within the genomes and uncover regions of local syntenic inheritance. We used Oxford Dot Plot [91] v0.3.3 to identify orthologous genes and plot synteny via ribbon plots, particularly the https://github.com/conchoecia/odp/blob/main/scripts/odp_nway_rbh pipeline. We extracted protein sequences from the annotated chromosomes of each Testudine assembly using the AGAT [92] v1.0.0 command *agat_sp_extract_sequences.pl* and mapped the sequences against each other using the diamond [88] v2.1.9 blastp command with e-value cutoff of 1e-5. Syntenic protein alignments were only included in the next step if the same hit was found to be the best for each pairwise comparison (best reciprocal hits). To determine syntenic blocks, permutation tests were performed, with 10,000 bootstraps and only those syntenic blocks with FDR less than 0.05 included and plotted as distinct colours in the ribbon diagrams. We performed this analysis once using the sea-turtle genomes as input and once with one species per genus for all currently available chromosome-scale reference genomes with annotations on GenBank alongside those from this study (GCF_016161935.1, GCF_007399415.2, GCF_028017835.1, GCF_013100865.1, GCF_027887155.1, GCF_009764565.3 and GCF_015237465.2).

### 6.6. Phylogenetic analysis

To reconstruct the phylogenetic history of the Testudine clade, we built a tree based on the protein sequences of all reference genomes submitted to GenBank with a protein-coding annotation. For each genome, we reduced the gff files to contain only the longest isoform per gene using the AGAT [92], v1.0.0 command *agat_sp_keep_longest_isoform.pl* and then extracted the protein sequences for each gene using the command *agat_sp_extract_sequences.pl*. To find the single-copy orthologs, we used OrthoFinder [93] v2.5.5.2 using the amino-acid files as input. We then aligned the single-copy orthologs using MAFFT [94] v7.475, trimmed the resulting multi-alignment files using trimAL [95] v1.4.1 with argument -automated1, concatenated the trimmed alignments into supermatrix containing all aligned sequences and constructed a phylogenetic tree using IQtree [96] v2.2.5 with 1,000 bootstraps (-B 1000). To further estimate the branching points in the tree, we took upper- and lower-bound divergence time estimates from timetree.org for all internal nodes and used these as calibration times for MCMCtree [97] paml v4.10.7 using the JC69 model. A full list of commands can be found in the script “create_tree.sh”.

### 6.7. Genome-wide diversity and divergence

Aiming to explore the genome-wide patterns of genetic diversity in the sea turtle clade, we performed the SNP calling for the seven sea turtle specie s using the jATG pipeline (https://github.com/diegomics/jATG/tree/devel). First, We mapped PacBio HiFi reads for the five genomes generated in this work against its own reference genome using minimap2 [84], v2.26, and mapped Illumina 10x reads for *D. coriacea* and *C. mydas* using bwa-mem2 [98] v2.2.1. Following the mapping, we removed PCR duplicates from BAM files using MarkDuplicates from GATK [99] v4.6. We performed variant calling using GATK v4.6 HaplotypeCaller and GenotypeGVCF. We then filtered the resulting GVCF using BCFtools [100] following GATK’s recommended parameter thresholds [99], removing low mapping quality positions (MQ>30), SNPs with depth lower than 8 and greater than 2× the average coverage, and keeping only biallelic positions. We also filtered small scaffolds, keeping only the 28 chromosomes for the subsequent analysis. Additionally, we excluded SNPs located in masked regions from subsequent analyses, identified by masking the genome with Dfam TE Tools v1.85 using RepeatModeler [101] and RepeatMasker [102]. We converted all filtered genotypes to missing data, producing a base-pair resolution gVCF file.

This filtered gVCF was used for genome-wide heterozygosity assessment and runs of homozygosity (RoH) analysis using Darwindow [103]. This tool enables the visualisation of heterozygosity and RoH along the scaffolds, providing a clear visual assessment of the accuracy of the RoH calls. We calculated heterozygosity based on a sliding-window approach with non-overlapping windows of 50 kb, without applying a filter for missing data. We identified RoHs using a heterozygosity threshold calculated from the average genome-wide heterozygosity of each species. A window was considered to have low heterozygosity if its value fell below one-fifth of the mean heterozygosity. The minimum length of a RoH was set to 500 kb, composed of at least 10 adjacent windows of 50 kb. The maximum proportion of missing data per window was 0.7. The inbreeding level (FRoH) was calculated as the proportion of the genome marked as RoH.

We calculated gene density using a custom Python script (GeneDensityCalculation.py) that counts the number of genes per Mb across the genome. We estimated pairwise genetic distances using a window-based approach, leveraging genome alignments generated with Progressive Cactus [104] v2.9. We used the halSnps pipeline to identify interspecific single variants, and the halAlignmentDepth pipeline to define 10 kb windows of aligned regions across the genome. We defined genetic distance in each window as the ratio of interspecific single variants per 10 kb. We then identified hotspots of genetic divergence, diversity and gene density by screening these metrics along the chromosomes and targeting windows where heterozygosity was higher than four times the chromosome mean and genetic distance exceeded twice the chromosome mean.

### 6.8. Demographic analysis

We inferred the demographic histories of the five sea turtle species whose genomes were assembled in this study using the Pairwise Sequentially Markovian Coalescent (PSMC) model [44]. We first extracted the consensus sequence from the filtered gVCF files generated above with BCFtools, then converted the resulting consensus fasta file into the PSMC input format using fq2psmcfa. We ran PSMC using default parameters: -N25 -t15 -r5 -p “4+25*2+4+6”, scaling the output assuming a mutation rate (μ) of 1.2 × 10⁻⁸ per site per generation and a generation time of 30 years. Given the uncertainty in generation time estimates across species and the variability reported in the literature for each, we selected a generation time of 30 years as an approximate midpoint of reported values. This choice provides a reasonable and biologically plausible basis for our analyses, as previously tested by Bentley et al. 2023 [34].

### 6.9 Hotspot gene family annotation

To further refine the annotation of gene families in the identified hotspot regions, we widened the search to include more functional databases available in InterProScan. Using the *D. coriacea* RefSeq annotation, we extracted the amino-acid sequence of the longest isoform for each gene using *agat_sp_keep_longest_isoform.pl* and the protein sequence using *agat_sp_extract_sequences.pl* as above and used these protein sequences as input to InterProScan (InterPro v102.0), using the Pfam [105], PRINTS [106], SUPERFAMILY [107], PANTHER [108], Gene3D [109], FunFam [110] and SMART [111] databases. To annotate genes belonging to multi-copy gene families, we classified genes as “MHC”, “Immunology-related”, “G-Protein Coupled Receptor” (GPCR), “Olfactory Receptor” or “Zinc-Finger” following terminology described in Table S9. Genes that did not fall into any of these multi-copy gene families were classified as “Other”. We then tested the enrichment of these multi-copy gene families against all other protein-coding genes annotated via Fisher’s exact test, followed by Benjamini-Hochberg correction of p-values to account for multiple testing. Full R script is available in the file “enrichment_test.R”.

## 7. Data Availability

Sequencing data, genome assemblies and annotations are available via the European Nucleotide Archive (ENA) or National Centre for Biotechnology Information (NCBI) under the following umbrella BioProjects: *Caretta caretta* PRJNA1212178, *Eretmochelys imbricata* PRJNA1212183, *Lepidochelys kempii* PRJNA1212180, *Lepidochelys olivacea* PRJNA1212179 and *Natator depressus* PRJNA1212185. Workflows used to generate genome assemblies are published on WorkflowHub under the following collection: https://workflowhub.eu/collections/10 and including the BioNano scaffolding workflow from the Vertebrate Genomes Project https://workflowhub.eu/workflows/643. To perform read mapping and SNP calling as well as calculating runs of homozygosity, we used the jATG pipeline available on GitLab: https://github.com/diegomics/jATG/tree/devel. Custom scripts written are published in gitlab: https://git.imp.fuberlin.de/begendiv/sea_turtlegenomes. Commands run to generate protein-coding annotations are available in the script “annotation_commands.sh”, custom script to calculate gene density is in the python file “geneDensityCalculation.py”.

## Declarations

*N. depressus* blood was collected under permit (TFA 2019-0174-2), animal ethics committee approval (2019-12B), and shipped under CITES permit (AU94). *L. kempii* blood and tissue samples were collected under USFWS Permit ES69328D. *E. imbricata* blood was collected under USFWS Permit TE-72088A-3. *C. caretta* and *L. olivacea* blood was collected under USFWS Permit TE86356B-2 (to Sea World), and *C. caretta* embryo tissue samples were collected under Florida Fish and Wildlife Conservation Commission Marine Turtle Permit 073 (FWC-MTP-073).

## Authors’ contributions

Conceptualization’s, LMK, OB, PHD, BPB

Data Curation: DDP, TB, JB

Formal Analysis: DDP, LSA, TB

Funding Acquisition: CJM, LMK, PHD, BPB, OB

Investigation: DDP, LSA, TB

Methodology: DDP, LSA, TB, CJM, LMK, PHD, BPB, OB, JB, CW, NJ, TT, BO’T, PT

Project Administration: CJM, LMK

Software: DDP, LSA, TB

Resources: SDW, GC, AK, DE, ELC, OB, PHD, CJM, LMK

Supervision: CJM, LMK, PHD, OB, EDJ

Validation: DDP, LSA, TB, CJM, LMK, PHD, BPB, OB

Visualisation: DDP, LSA, TB

Writing - Original Draft: LSA, TB

Writing - Review & Editing: All

## Acknowledgements

The authors would like to thank the HPC Service of FUB-IT, Freie Universität Berlin, for computing time [112], Camryn Allen, Shreya Banerjee, Jamie Adkins, Alexandria Mena, Andra Kurtz, Claudia Cedillo, Itzel Sifuentes-Romero, Jeanette Wyneken and The Rescue and Rehabilitation Department and Animal Health Department of New England Aquarium, especially Charlie Innis, for assistance with sample collection, and the Revive & Restore team for support with project planning.

