## Supplementary_Material for "Haplotype-resolved reference genomes of the sea turtle clade unveil ultra-syntenic genomes with hotspots of divergence"

### Supplementary Figures

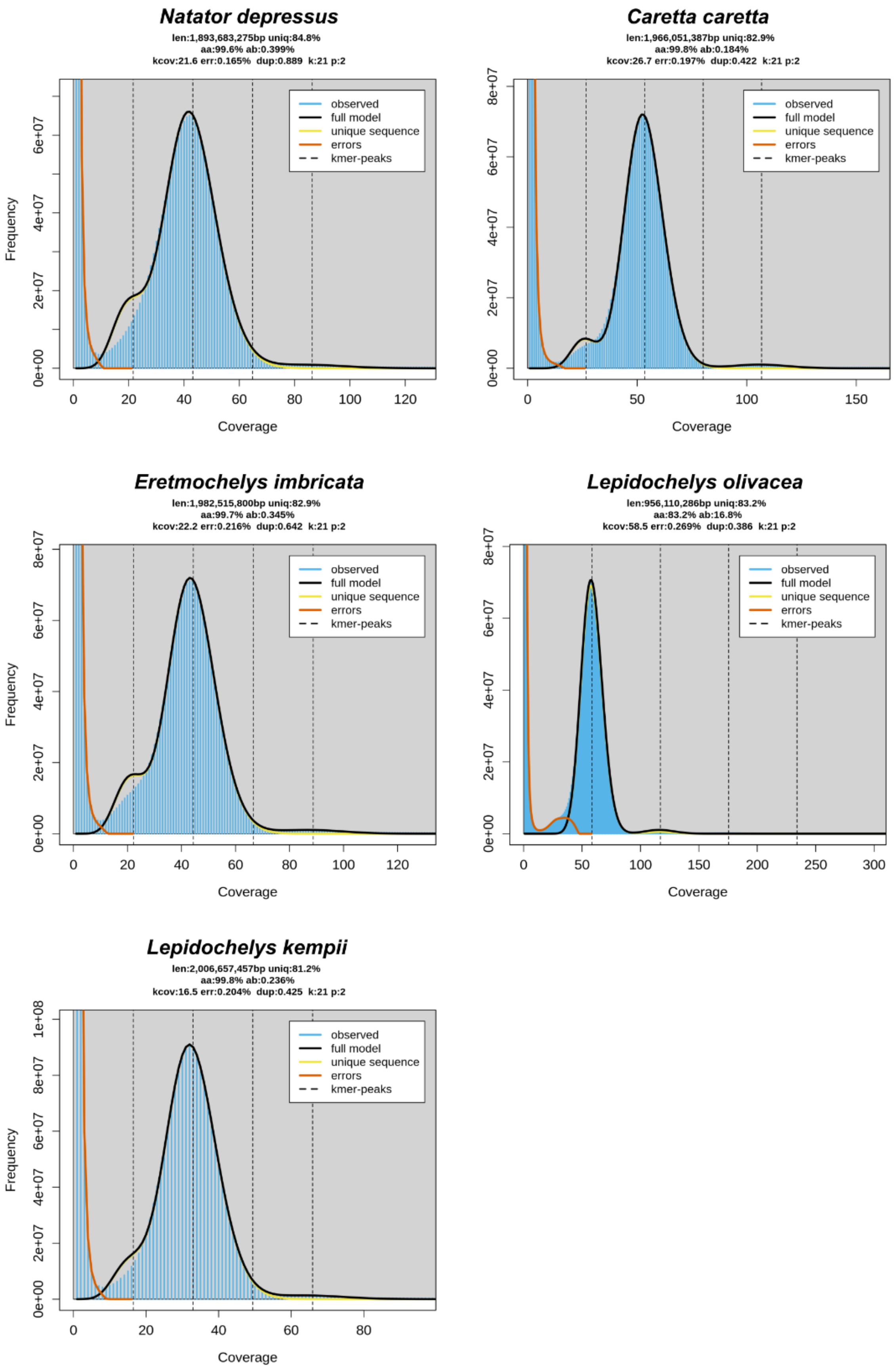

Natator depressus
len:1,893,683,275bp uniq:84.8%
aa:99.6% ab:0.399%
kcov:21.6 err:0.165% dup:0.889 k:21 p:2
Caretta caretta
len:1,966,051,387bp uniq:82.9%
aa:99.8% ab:0.184%
kcov:26.7 err:0.197% dup:0.422 k:21 p:2
Eretmochelys imbricata
len:1,982,515,800bp uniq:82.9%
aa:99.7% ab:0.345%
kcov:22.2 err:0.216% dup:0.642 k:21 p:2
Lepidochelys olivacea
len:956,110,286bp uniq:83.2%
aa:83.2% ab:16.8%
kcov:58.5 err:0.269% dup:0.386 k:21 p:2
Lepidochelys kempii
len:2,006,657,457bp uniq:81.2%
aa:99.8% ab:0.236%
kcov:16.5 err:0.204% dup:0.425 k:21 p:2
observed
full model
unique sequence
errors
kmer-peaks
Frequency
Coverage

**Sup. Fig. 1** K-mer based genome profiling for the sequenced sea-turtles. Shown are the distributions of 31-mers in each PacBio HiFi dataset for the sequenced individuals. Overlaid are the genome profile models as calculated by GenomeScope, giving indications of genome size, heterozygosity (ab), non-repetitive content of the genome (uniq) as well as estimates of the heterozygous genome coverage (kcov), read error (err) and duplication rate (dup)

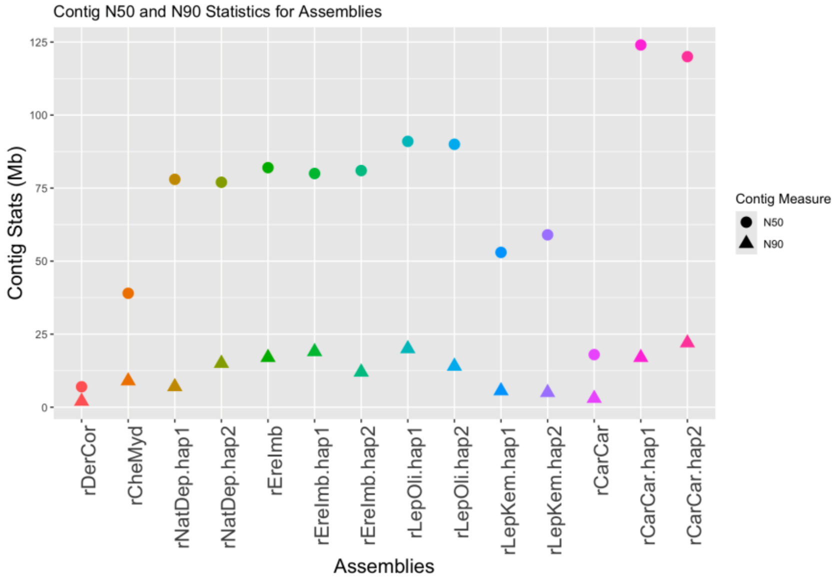


**Sup. Fig. 2** Genome contiguity statistics of sea turtle genomes. Shown are the Contig N50 and N90 values for all available sea-turtle genomes. N50 values are shown as circles and N90 values as triangles. Included are the 10 haplotype-separated assemblies from this study, PacBio CLR-based assemblies for *Dermochelys* and *Chelonia*, an ONT-based assembly for *Caretta* and one PacBio HiFi assembly for *Eretmochelys*.

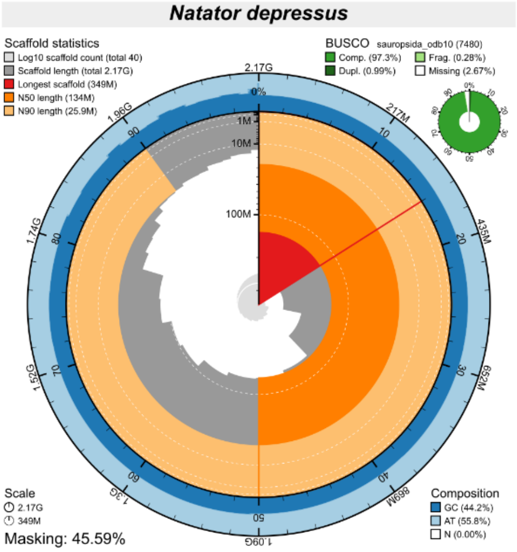
Natator depressus
Scaffold statistics
Log10 scaffold count (total 40)
Scaffold length (total 2.17G)
Longest scaffold (349M)
N50 length (134M)
N90 length (25.9M)
BUSCO sauropsida_odb10 (7480)
Comp. (97.3%)
Frag. (0.28%)
Dupl. (0.99%)
Missing (2.67%)
Scale
2.17G
349M
Masking: 45.59%
Composition
GC (44.2%)
AT (55.8%)
N (0.00%)

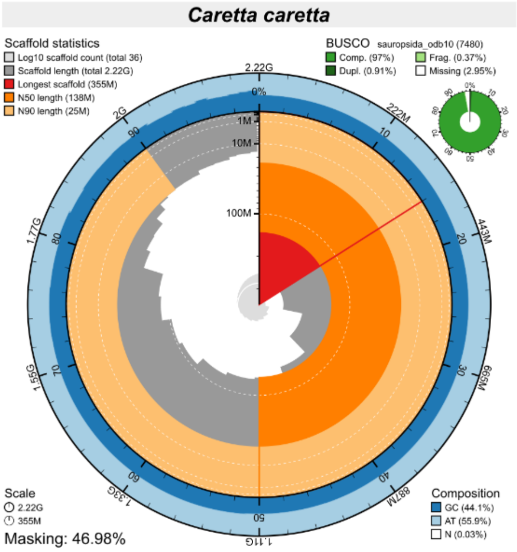
Caretta caretta
Scaffold statistics
Log10 scaffold count (total 36)
Scaffold length (total 2.22G)
Longest scaffold (355M)
N50 length (138M)
N90 length (25M)
BUSCO sauropsida_odb10 (7480)
Comp. (97%)
Frag. (0.37%)
Dupl. (0.91%)
Missing (2.95%)
Scale
2.22G
355M
Masking: 46.98%
Composition
GC (44.1%)
AT (55.9%)
N (0.03%)

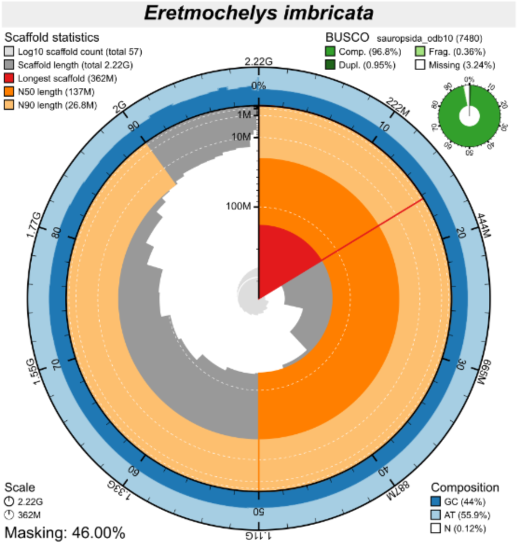
Eretmochelys imbricata
Scaffold statistics
Log10 scaffold count (total 57)
Scaffold length (total 2.22G)
Longest scaffold (362M)
N50 length (137M)
N90 length (26.8M)
BUSCO sauropsida_odb10 (7480)
Comp. (96.8%)
Frag. (0.36%)
Dupl. (0.95%)
Missing (3.24%)
Scale
2.22G
362M
Masking: 46.00%
Composition
GC (44%)
AT (55.9%)
N (0.12%)

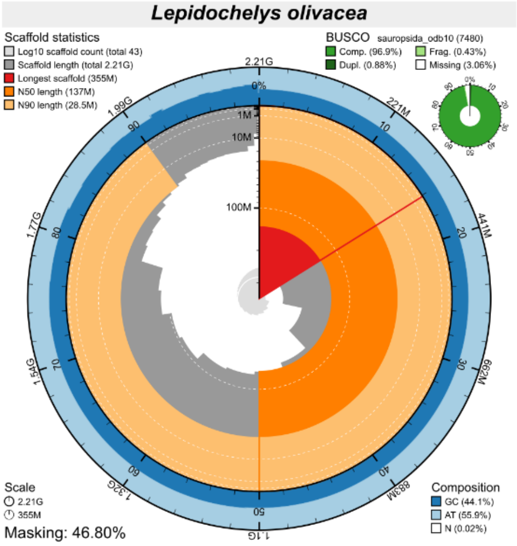
Lepidochelys olivacea
Scaffold statistics
Log10 scaffold count (total 43)
Scaffold length (total 2.21G)
Longest scaffold (355M)
N50 length (137M)
N90 length (28.5M)
BUSCO sauropsida_odb10 (7480)
Comp. (96.9%)
Frag. (0.43%)
Dupl. (0.88%)
Missing (3.06%)
Scale
2.21G
355M
Masking: 46.80%
Composition
GC (44.1%)
AT (55.9%)
N (0.02%)

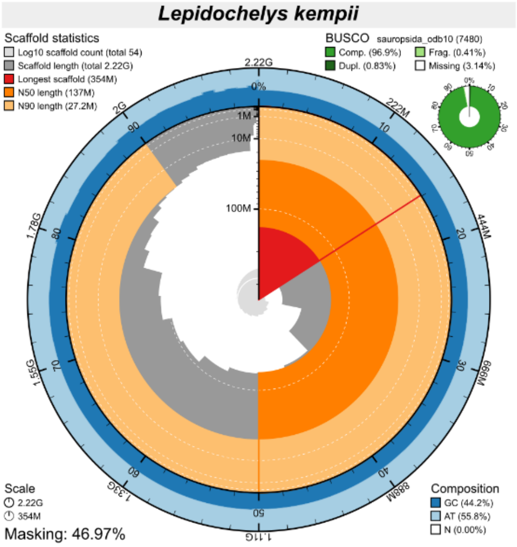
Lepidochelys kempii
Scaffold statistics
Log10 scaffold count (total 54)
Scaffold length (total 2.22G)
Longest scaffold (354M)
N50 length (137M)
N90 length (27.2M)
BUSCO sauropsida_odb10 (7480)
Comp. (96.9%)
Frag. (0.41%)
Dupl. (0.83%)
Missing (3.14%)
Scale
2.22G
354M
Masking: 46.97%
Composition
GC (44.2%)
AT (55.8%)
N (0.00%)

**Sup. Fig. 3** Snail plots summary of assembly statistics. The main plot is divided into 1,000 size-ordered bins around the circumference, with each bin representing 0.1% of the total assembly. The distribution of sequence lengths is shown in dark grey, with the plot radius scaled to the longest sequence present in the assembly (shown in red). Orange and pale-orange arcs show the scaffold N50 and N90 sequence lengths, respectively. The pale grey spiral shows the cumulative sequence count on a log-scale, with white scale lines showing successive orders of magnitude. The blue and pale-blue area around the outside of the plot shows the distribution of GC, AT, and N percentages in the same bins as the inner plot. A summary of complete, fragmented, duplicated, and missing BUSCO genes found in the assembled genome from the Sauropsida database (odb10) is shown in the top right.

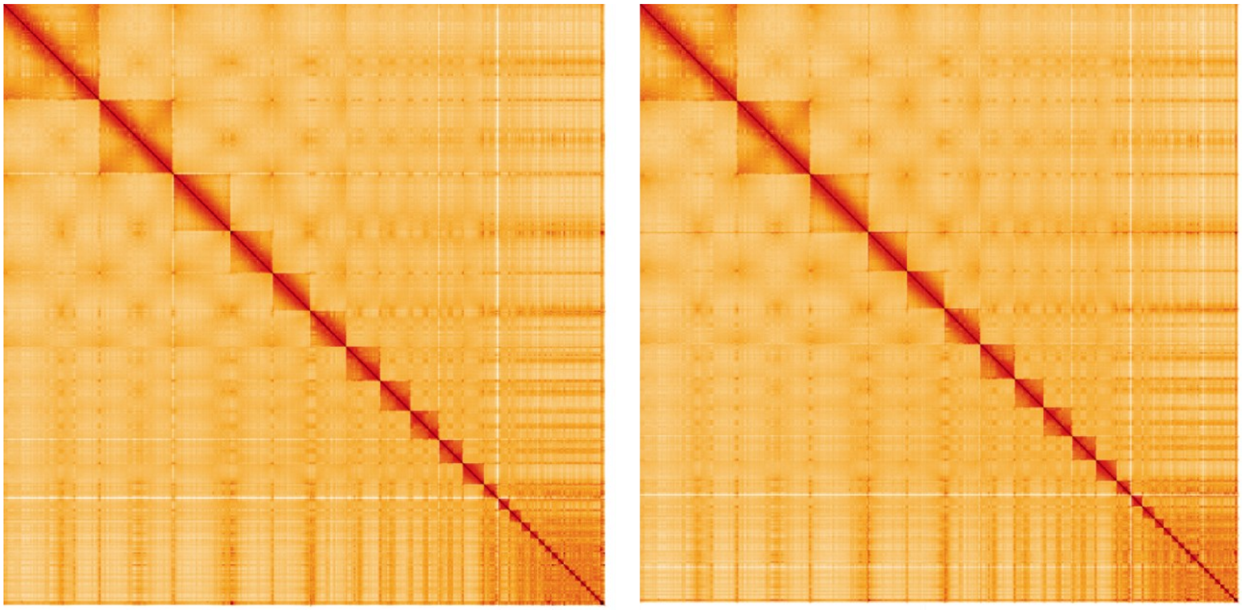

*Caretta caretta* Hi-C interactions

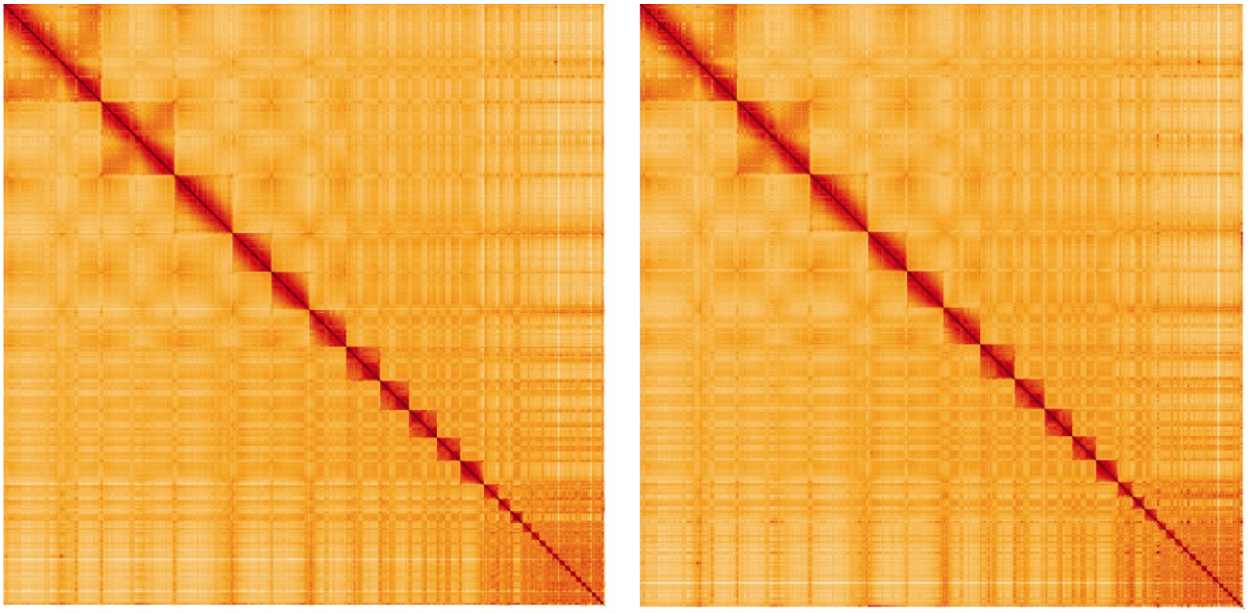

*Eretmochelys imbricata* Hi-C interactions

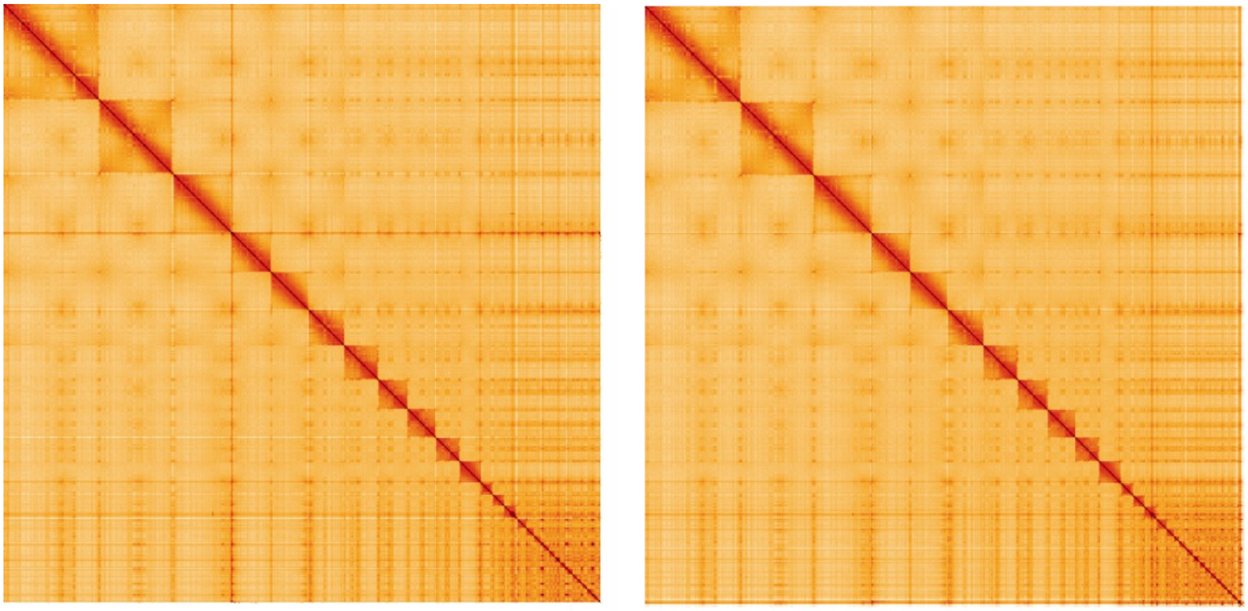

*Lepidochelys olivacea* Hi-C interactions

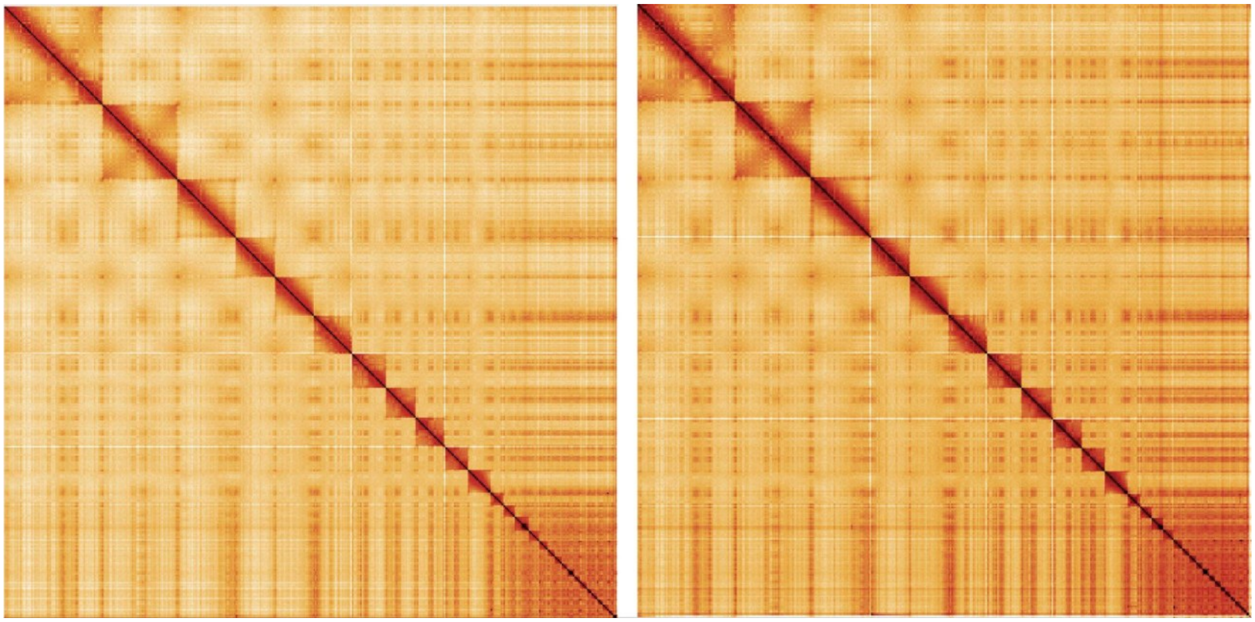

*Lepidochelys kempii* Hi-C interactions

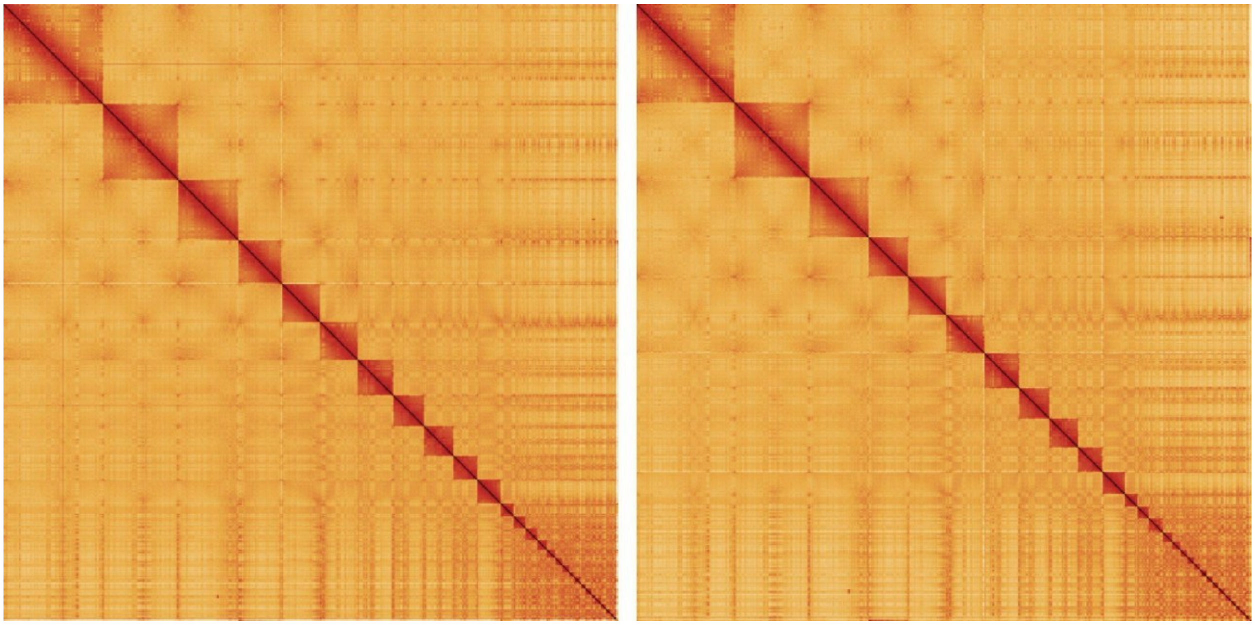

*Natator depressus* Hi-C interactions

**Sup. Fig. 4** 3-dimensional conformational arrangement of the sea-turtle genomes as evaluated by Hi-C. The x- and y-axes show the coordinates of the respective genome and each detected contact in the genome is coloured with increasing intensity from white to red. The red diagonal shows the self-interactions of each position with itself and close vicinity.

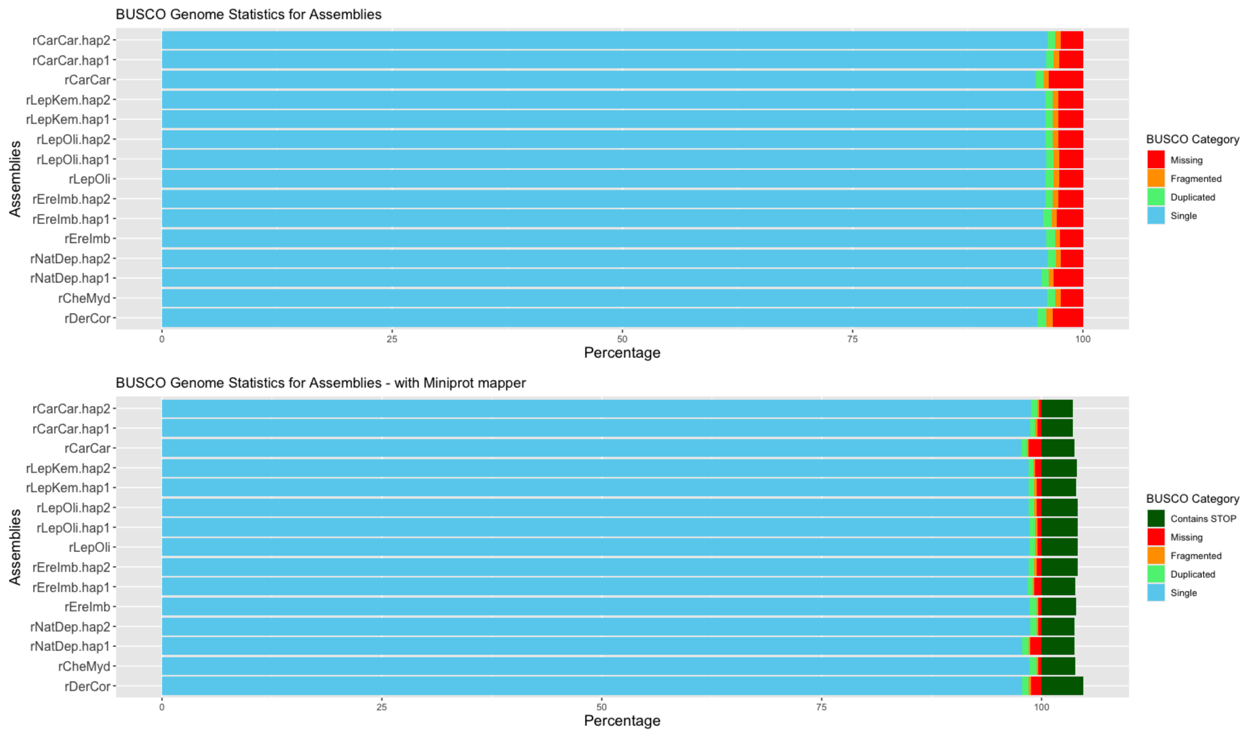


**Sup. Fig. 5** Gene completeness of sea-turtle genomes. Shown are the percentage detected single copy orthologs calculated via BUSCO using the Sauropsida database. Scores are calculated based on detected mappings of ortholog sequences using Metaeuk (above) and Miniprot (below). Note the Miniprot mode of BUSCO also gives estimates of BUSCO genes potentially containing early STOP codons.

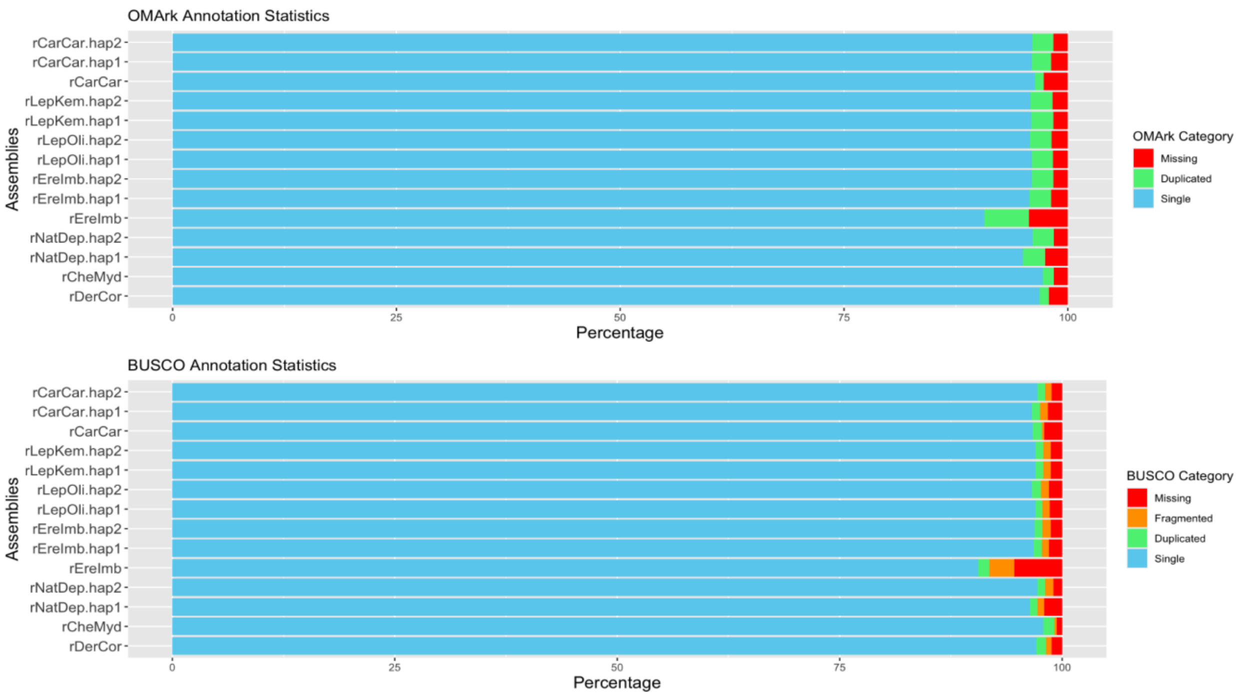


**Sup. Fig. 6** Gene completeness of sea-turtle protein-coding annotations. Shown are the percentage Sauropsida Hierarchical Orthology Groups (above) and single copy orthologs (below) as calculated by OMArk and BUSCO, respectively, from the annotated protein sequences in each sea-turtle genome.

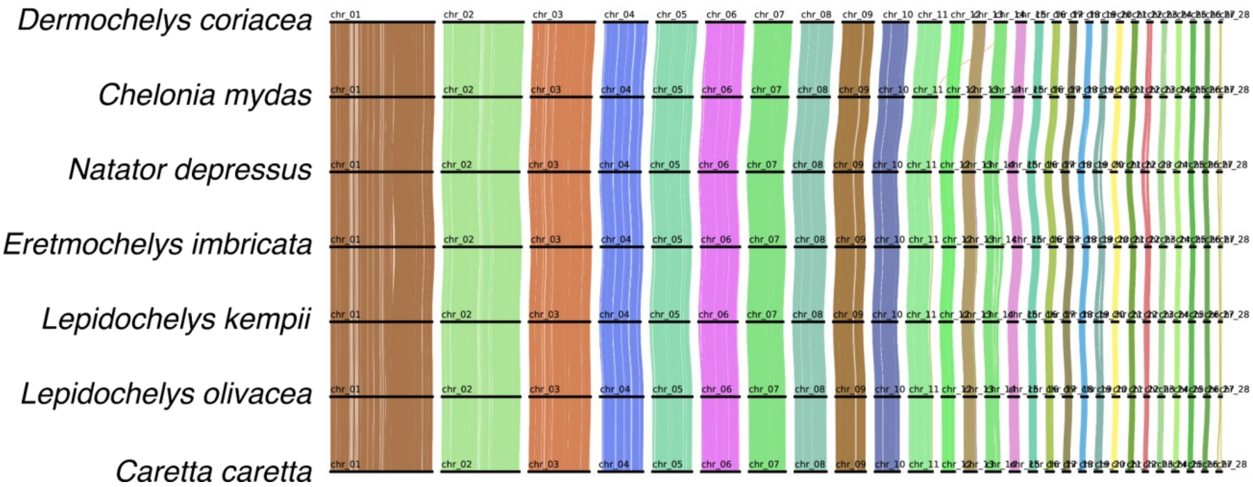


**Sup. Fig. 7** Genome-wide gene-synteny plots for sea turtles. Each line represents a best-reciprocal-hit protein match between annotated genes in each consecutive genome. Lines are coloured based on co-localisation across all 7 genomes determined by Fisher's Exact Test. Chromosomes are ordered based on synteny to *Dermochelys coriacea* genome.

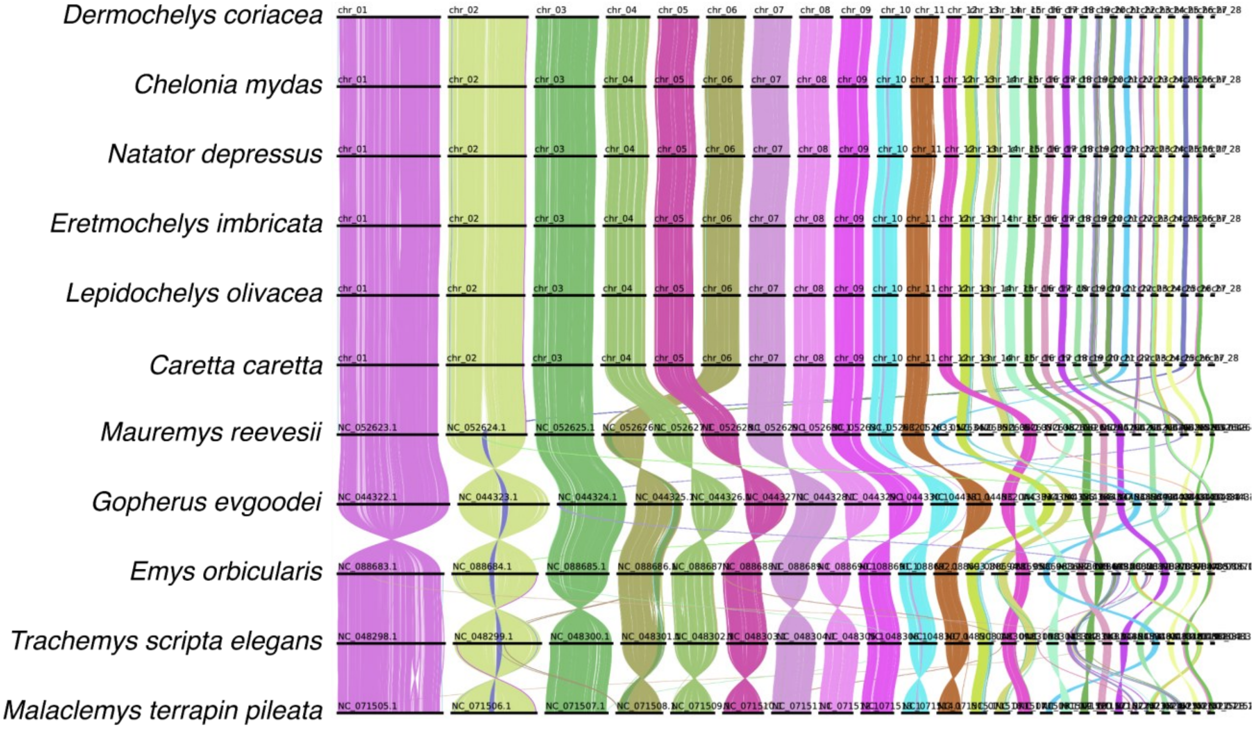


**Sup. Fig. 8** Genome-wide gene-synteny plots across testudines. Each line represents a best-reciprocal-hit protein match between annotated genes in each consecutive genome. Lines are coloured based on co-localisation across all 11 genomes determined by Fisher's Exact Test. Chromosomes are ordered based on the assigned chromosome name in GenBank.

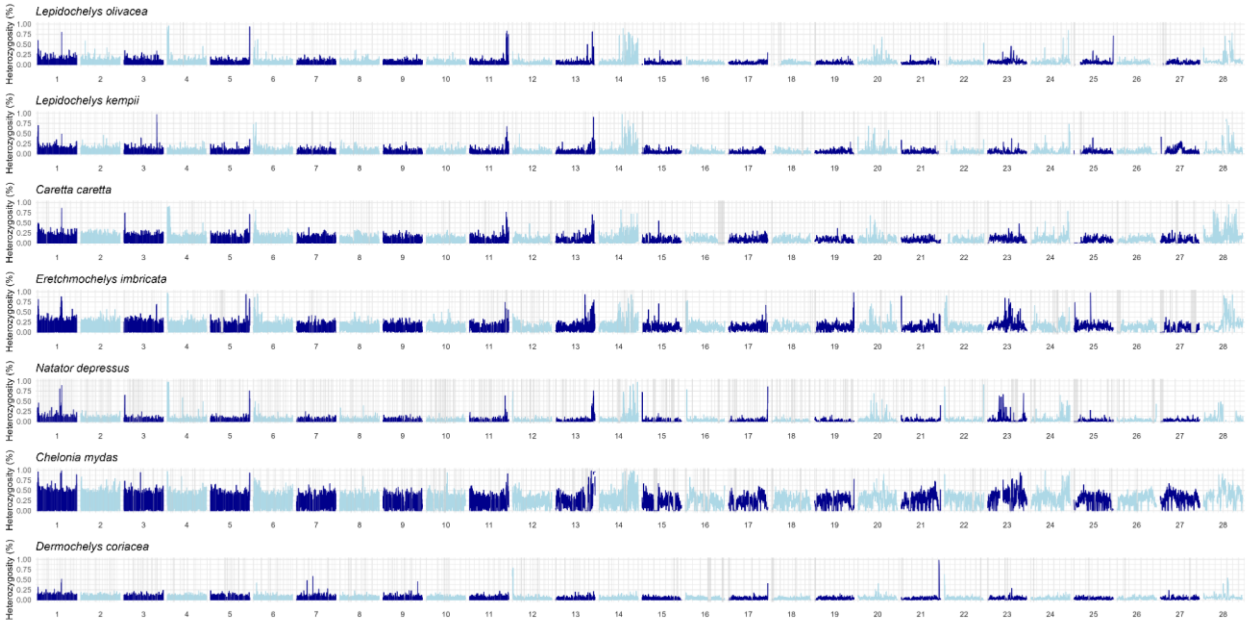


**Sup. Fig. 9** Variation of genome-wide heterozygosity (He) across the 28 chromosomes for the seven sea turtle species, depicting He of non-overlapping 50 kb windows with RoH segments highlighted in gray. Chromosomes are represented with equal widths, therefore RoHs sizes are not proportional among them as depicted in the figure.

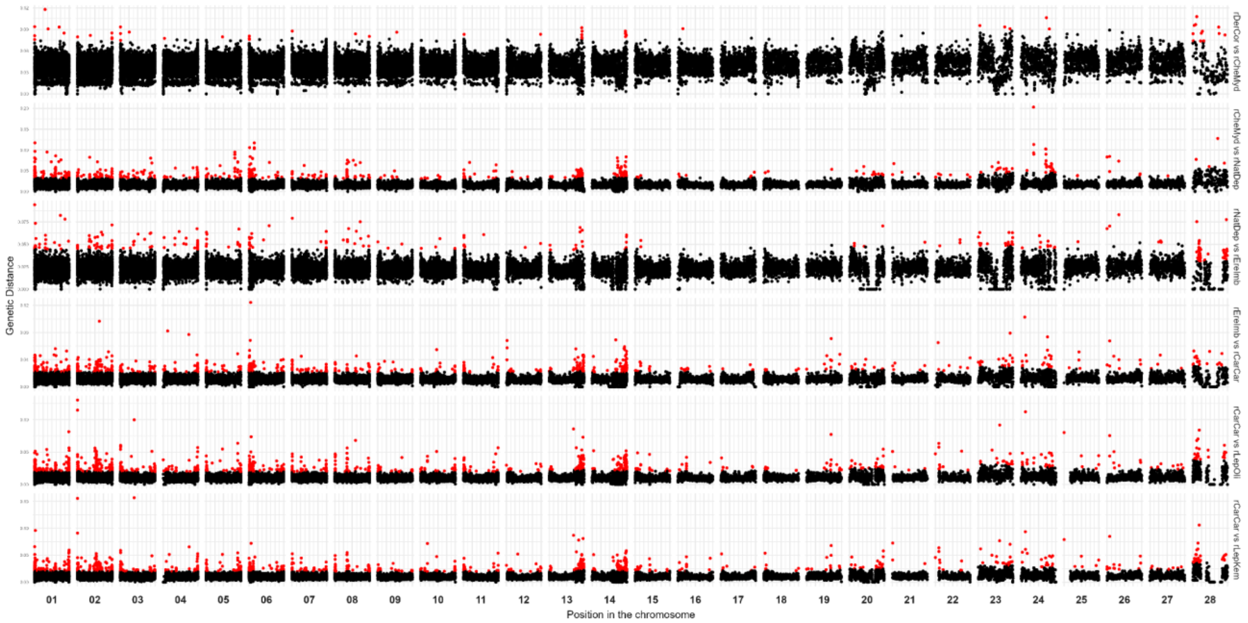


**Sup. Fig. 10** Pairwise genetic distances between different sea turtle species along the 28 chromosomes. The first species listed in each comparison refers to the genome considered as the reference in the Cactus alignment. Genetic distance was calculated as the ratio of interspecific single variants per 30 kb window. Red dots represent windows with genetic distance greater than 2x the average. Chromosomes are shown with equal widths.

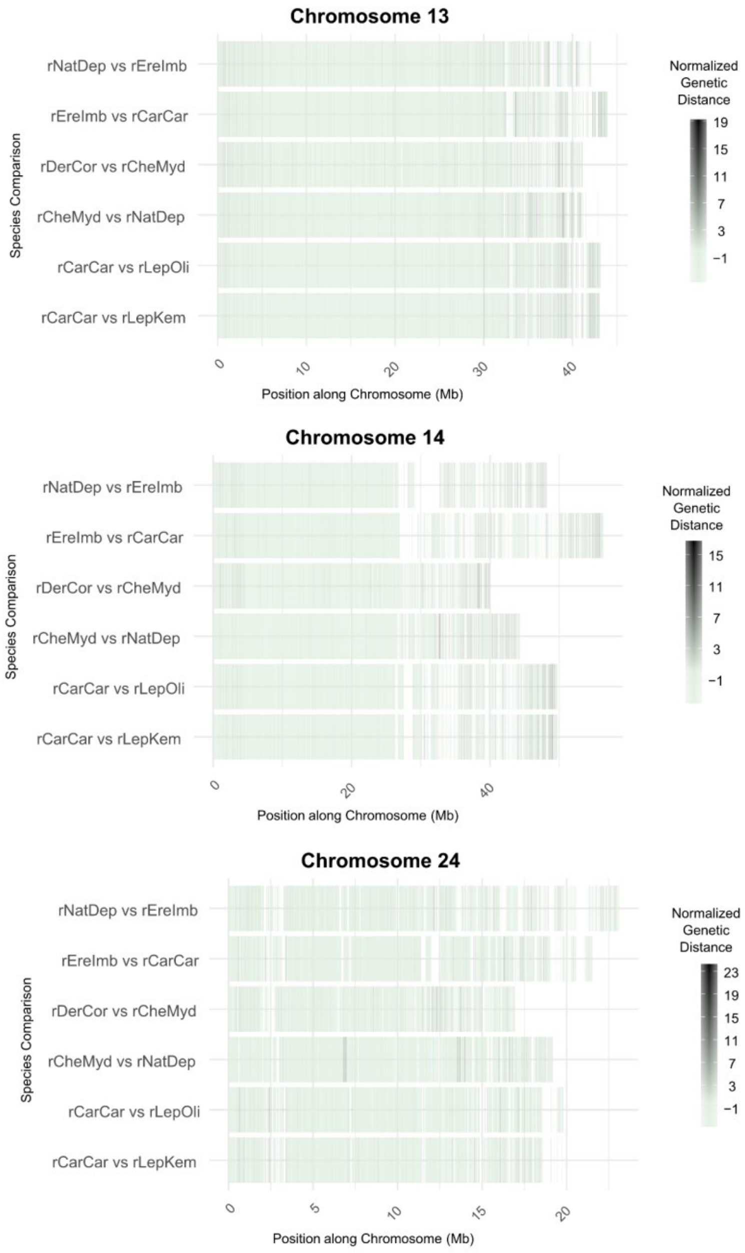


**Sup. Fig. 11** Pairwise genetic distances between different sea turtle species along the chromosomes 13, 14 and 24. The first species listed in each comparison refers to the genome considered as the reference in the Cactus alignment. Genetic distance was calculated as the ratio of interspecific single variants per 10 kb window, with normalization applied to highlight chromosomal hotspots rather than overall species divergence.

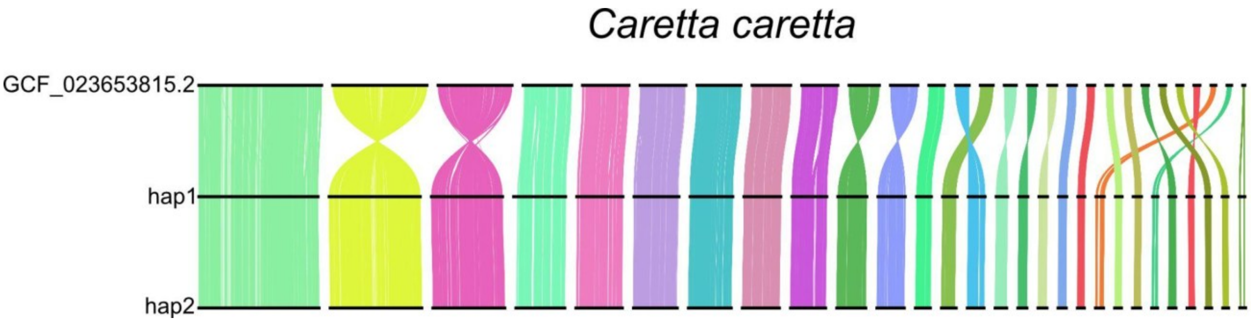


**Sup. Fig. 12** Ribbon plot showing the locations of syntenic genes between the previous available genome from *Caretta caretta* (top) and the two haplotype assemblies presented in this study

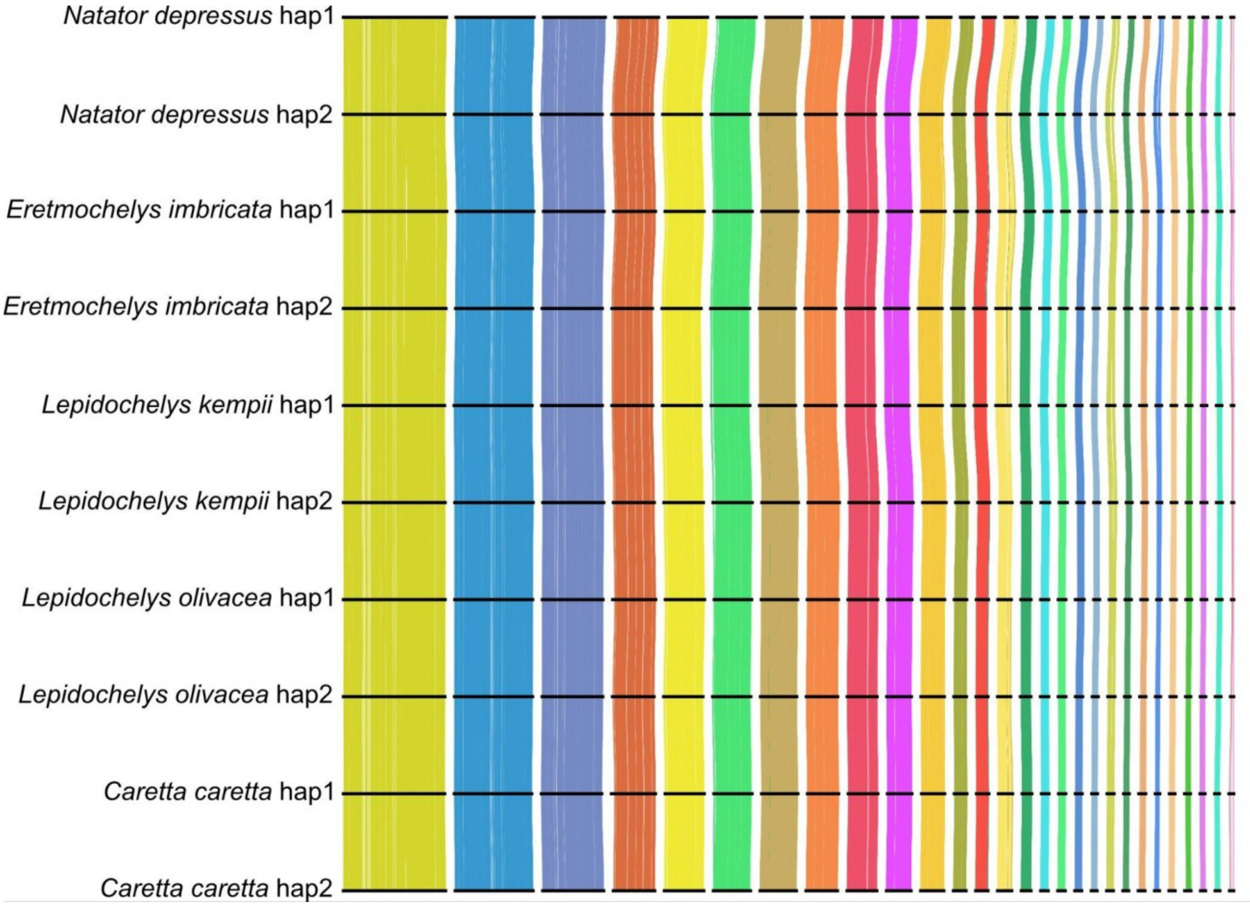


**Sup. Fig. 13** Ribbon plot showing the locations of syntenic genes between all as asemblies presented in this study

### Supplementary Tables

**Table S1** PacBio HiFi and OmniC Hi-C sequencing coverage after adapter and quality trimming.

| | HiFi | | Hi-C | | Bionano | |
|---|---|---|---|---|---|---|
| Asm ID | bp | x | bp | x | Gbp (≥150kbp) | x |
| rCarCar | 121,592,450,120 | 55 | 264,606,173,591 | 120 | 507.12 | 231 |
| rEreImb | 101,064,681,912 | 46 | 283,870,332,269 | 129 | 109.12 | 50 |
| rLepKem | 76,007,057,258 | 35 | 102,633,790,121 | 47 | - | - |
| rLepOli | 132,137,130,189 | 60 | 188,733,679,226 | 86 | 521.74 | 237 |
| rNatDep | 91,008,047,949 | 41 | 166,770,516,289 | 76 | - | - |

**Table S2** Assembly metrics for all the available sea turtle assemblies.
Excel format.

**Table S3** Statistics from protein-coding annotations.

| Genome | No. genes | Mean gene length (bp) | Mean transcript length (bp) | Mean no. exons | No. single-exon genes |
|---|---|---|---|---|---|
| **rCarCar.hap1** | 24,378 | 40,283 | 1,663 | 8.8 | 4,669 |
| **rCarCar.hap2** | 26,029 | 37,848 | 1,683 | 8.3 | 5,985 |
| **rEreImb.hap1** | 25,692 | 38,220 | 1,585 | 8.3 | 5,682 |
| **rEreImb.hap2** | 25,792 | 38,033 | 1,589 | 8.4 | 5,485 |
| **rLepKem.hap1** | 26,436 | 37,158 | 1,620 | 8.2 | 6,372 |
| **rLepKem.hap2** | 25,992 | 37,419 | 1,622 | 8.2 | 6,360 |
| **rLepOli.hap1** | 21,475 | 44,628 | 1,715 | 9.7 | 2,432 |
| **rLepOli.hap2** | 24,071 | 40,107 | 1,671 | 8.9 | 4,446 |
| **rNatDep.hap1** | 25,069 | 37,948 | 1,618 | 8.2 | 5,969 |
| **rNatDep.hap2** | 26,687 | 36,581 | 1,626 | 8.2 | 6,158 |

**Table S4** Divergence time estimates for all nodes of the phylogenetic tree shown in figure 3.

| Node | Mean divergence time (MYA) | 95% highest posterior density (HPD) interval (MYA) |
|---|---|---|
| 1. Cryptodira | 164 | 147, 182 |
| 2. Durocryptodira | 104 | 81.9, 122 |
| 3. (Geoemydidae + Testudinidae) / (Platysternidae + Emydidae) | 79.6 | 65.4, 94.5 |
| 4. Geoemydidae / Testudinidae | 57.1 | 52.2, 61.3 |
| 5. Platysternidae / Emydidae | 55 | 29.5, 83.6 |
| 6. *Chelonoidis abingdonii* / *Gopherus* sp. | 48.4 | 39.0, 58.1 |
| 7. Emydidae | 39.7 | 23.4, 56.8 |
| 8. *Gopherus flavomarginatus* / *Gopherus evgoodei* | 24.1 | 0.00087, 46.9 |
| 9. *Chrysemys picta* / (*Malaclemys terrapin* + *Trachemys scripta*) | 23.7 | 12.6, 34.4 |
| 10. *Terrapene triunguis* / *Emys orbicularis* | 23.2 | 11.8, 36.1 |
| 11. *Mauremys reevesii* / *Mauremys mutica* | 21.7 | 15.6, 27.9 |
| 12. *Malaclemys terrapin* / *Trachemys scripta* | 14.6 | 8.48, 21.8 |
| 13. Dermochelyidae / Cheloniidae | 75.4 | 49.4, 104 |
| 14. (*Natator depressus* + *Chelonia mydas*) / (*Eretmochelys imbricata* + *Caretta caretta* + *Lepidochelys* sp.) | 48 | 33.7, 61.8 |
| 15. *Natator depressus* / *Chelonia mydas* | 33.6 | 33.5, 33.8 |

| | | |
|---|---|---|
| 16. *Eretmochelys imbricata* / (*Caretta caretta* + *Lepidochelys* sp.) | 25.4 | 17.4, 31.9 |
| 17. *Caretta caretta* / *Lepidochelys* sp. | 18.6 | 13.3, 24.7 |
| 18. *Lepidochelys olivacea* / *Lepidochelys kempii* | 7.72 | 2.99,12.4 |

**Table S5 -** Heterozygosity, inbreeding level (FRoH) and gene density per chromosome for the seven sea turtle reference genomes. This data is provided in a separate file in Excel format.

**Table S6** Enrichment p-values for multi-copy gene families in hotspots of genetic diversity

| | Chr 13 | Chr 14 | Chr 24 |
|---|---|---|---|
| MHC | 0.3785960 | $8.542079\text{x}10^{-26}$ | 1 |
| Immunology | $4.126994\text{x}10^{-44}$ | $3.892572\text{x}10^{-7}$ | $7.524291\text{x}10^{-5}$ |
| GPCR | $2.215344\text{x}10^{-49}$ | $1.645942\text{x}10^{-3}$ | $5.917677\text{x}10^{-5}$ |
| Olfactory | $2.320270\text{x}10^{-81}$ | $2.233347\text{x}10^{-10}$ | 0.1254281 |
| Zinc-finger | $5.914689\text{x}10^{-3}$ | $1.836372\text{x}10^{-54}$ | 0.8770295 |

**Table S7** Benjamini-Hochberg-corrected p-values for multi-copy gene families in hotspots of genetic diversity

| | Chr 13 | Chr 14 | Chr 24 |
|---|---|---|---|
| MHC | 0.4368415 | $2.562624\text{x}10^{-25}$ | 1 |
| Immunology | $1.547623\text{x}10^{-43}$ | $8.341225\text{x}10^{-7}$ | $1.254048\text{x}10^{-4}$ |
| GPCR | $1.107672\text{x}10^{-48}$ | $2.468912\text{x}10^{-3}$ | $1.109564\text{x}10^{-4}$ |
| Olfactory | $3.480405\text{x}10^{-80}$ | $5.583367\text{x}10^{-10}$ | 0.1567851 |
| Zinc-finger | $8.065485\text{x}10^{-3}$ | $1.377279\text{x}10^{-53}$ | 0.9396744 |

**Table S8** Weights used to combine transcriptome evidences into a single gene-model set

| Type | Evidence | Weight |
|---|---|---|
| ABINITIO_PREDICTION | Helixer | 1 |
| OTHER_PREDICTION | stringtie_rnaseq | 2 |
| OTHER_PREDICTION | stringtie_isoseq | 3 |
| OTHER_PREDICTION | toga_rMalTer | 4 |
| OTHER_PREDICTION | toga_rCheMyd | 5 |
| PROTEIN | rDerCor_miniprot | 6 |
| PROTEIN | rCheMyd_miniprot | 7 |

**Table S9** Database and terminology used for classifying genes based on their functional annotation.

| Database | Terms used for classifying genes as "MHC" |
|---|---|
| Pfam | "MHC class II, alpha chain, N-terminal" |
| PRINTS | "MHC class I alpha chain, alpha1 alpha2 domains" |
| PANTHER | "Major Histocompatibility Complex/Immunoglobulin" & "Antigen-presenting and immune regulatory MHC class I-related" |
| Gene3D | "MHC class II, alpha/beta chain, N-terminal" |
| FunFam | "Major histocompatibility complex, class I-related protein", "H-2 class I histocompatibility antigen, alpha chain" & "HLA class II histocompatibility antigen, DRB1-1 beta chain" |
| SMART | "MHC class II, alpha chain, N-terminal" |
| **Database** | **Terms used for classifying genes as "Immunology-related"** |
| Pfam | "CD80-like C2-set immunoglobulin domain", "Immunoglobulin I-set domain", "Immunoglobulin V-set domain", "Immunoglobulin domain", "Immunoreceptor tyrosine-based activation motif", "Immunoglobulin-like beta-sandwich domain" & "Phosphorylated immunoreceptor signalling ITAM" |

| | |
|---|---|
| SUPERFAMILY | “Immunoglobulin-like domain superfamily” |
| PANTHER | “Immunoglobulin superfamily BTN/MOG domain-containing protein”, “Immunoglobulin superfamily CEA-related”, “Immunoglobulin-like domain-containing protein LISCH7”, “SIALIC ACID BINDING IMMUNOGLOBULIN-LIKE LECTIN” & “TRANSMEMBRANE AND IMMUNOGLOBULIN DOMAIN-CONTAINING PROTEIN” |
| Gene3D | “Immunoglobulin-like fold” |
| SMART | “Immunoglobulin domain subtype”, “Immunoglobulin subtype 2”, “Immunoglobulin V-set domain” & “ Phosphorylated immunoreceptor signalling ITAM” |
| **Database** | **Terms used for classifying genes as “G-Protein Coupled Receptor (GPCR)”** |
| Pfam | “G protein-coupled receptor, rhodopsin-like” |
| PRINTS | “G protein-coupled receptor 40-related receptor” & “G protein-coupled receptor, rhodopsin-like” |
| SUPERFAMILY | “Family A G protein-coupled receptor-like” |
| FunFam | “G-protein coupled receptor” |
| **Database** | **Terms used for classifying genes as “Olfactory Receptor”** |
| Pfam | “Olfactory receptor” |
| PRINTS | “Olfactory receptor” |
| PANTHER | “Human Olfactory Receptors”, “Human and Mouse Olfactory Receptor”, “Olfactory G-protein coupled receptor”, “Olfactory receptor subfamily 6C-like” & “OLFACTORY RECEPTOR” |
| FunFam | “Olfactory receptor” |
| **Database** | **Terms used for classifying genes as “Zinc-Finger”** |
| Pfam | “B-box-type zinc finger”, ”Zinc finger C2H2-type”, ”Zinc finger, BED-type”, ”Zinc finger, C3HC4 RING-type”, ”Zinc finger, CCCH-type”, ”Zinc finger, PHD-finger”, ”Zinc finger, RING-type”, ”Zinc finger, RING-type, eukaryotic”, ”Zinc finger, TFIIS-type”, “Transcription factor S-II (TFIIS) Zinc finger”, “Zinc-finger double domain”, “zinc finger of C3HC4-type, RING” & “C2H2-type zinc finger” |

| | |
|---|---|
| PRINTS | “Zinc finger, B-box, chordata” |
| SUPERFAMILY | “B-box zinc-binding domain”, “CCCH zinc finger”, “FYVE/PHD zinc finger”, “beta-beta-alpha zinc fingers” |
| PANTHER | “ZINC FINGER PROTEIN”, “ZINC FINGER AND BTB DOMAIN-CONTAINING”, “ZINC FINGER AND SCAN DOMAIN-CONTAINING”, “ZNF44 PROTEIN”, “Zinc finger BED domain-containing protein”, “C2H2-type zinc-finger domain-containing protein” & “Krueppel C2H2-type Zinc Finger” |
| Gene3D | “Zinc finger, RING/FYVE/PHD-type” & “Classic Zinc Finger” |
| FunFam | “zinc finger protein”, “zinc finger and SCAN domain-containing protein”, “Zinc finger with KRAB and SCAN domains”, “zinc finger family member”, “KRAB zinc finger”, “Zinc finger 45-like” |
| SMART | “B-box-type zinc finger”, “Zinc finger C2H2-type”, “Zinc finger, CCCH-type”, “Zinc finger, PHD-type”, “Zinc finger, RING-type” & “Zinc finger, TFIIS-type” |
